# Torque-generating units of the bacterial flagellar motor are rotary motors

**DOI:** 10.1101/2025.10.28.684902

**Authors:** Basarab G. Hosu, Alina M. Vrabioiu, Aravinthan D.T. Samuel

## Abstract

*E. coli* swims using helical flagellar filaments driven at their base by a rotary motor. Torque-generating ‘stator’ units drive the bacterial flagellar motor (BFM) by transmitting mechanical power to a cytoplasmic ‘rotor’, the C-ring. Each stator unit is a proton-conducting heteromer. A central dimer of two MotB proteins anchor to the cell wall. A surrounding pentamer of five MotA proteins transmit mechanical power to the C-ring. This asymmetrical 5:2 structure is consistent with rotation as the mechanism of torque generation. Here, we test the hypothesis that the MotA_5_MotB_2_ stator units are rotary motors themselves and interact with the rotor like intermeshed gearwheels, where rotation of the C-ring is directly coupled to MotA_5_ rotation around the MotB_2_. We used *in vivo* polarized photo-bleaching microscopy. When a subset of fluorescent domains inside a multimer is rapidly photo-bleached by a strong pulse of polarized light, the induced polarization-dependent fluorescence of unbleached domains becomes a reporter of angular orientation. We applied polarized photo-bleaching microscopy to tethered cells rotating by single flagellar motors. We probed fluorescently-labeled MotA pentamer and MotB dimer calibrated to motor rotation. The MotB dimer rotates at the same angular speed as the cell body, consistent with its anchor to the cell wall. The MotA pentamer rotates ∼6.2x faster than the flagellar motor, revealing the gear ratio between stator and rotor.

**Significance Statement:** Bacteria swim by rotating rigid helical flagellar filaments. Here, we find that the torque-generating unit that drives flagellar rotation is itself a rotary motor. Each torque-generating unit is a heteromeric macromolecular machine – a pentamer of MotA subunits that surround a dimer of proton-conducting MotB subunits. Torque is generated as the MotA spins around MotB. The MotA pentamer interacts with rotor of the flagellar motor in a manner resembling intermeshing gearwheels. The bacterial flagellar motor is driven by the first set of enmeshed gearwheels that has been described in any living cell.

---

The bacterial flagellar motor (BFM) is a proton-powered rotary engine. (1–3) Cryo-electron microscopy has revealed the structure of the rotor at the flagellar base (Fig. 1A) and the structure and stoichiometry of the torque-generating stator unit (Fig. 1B). (4–6) The rotor has a central rod with rings for each layer of the cell envelope. Outside the cell, the rod connects to the flagellar filament by a flexible hook. Inside the cell, the rod connects to the MS-ring, in turn attached to the C-ring, the wheel that is mechanically powered by surrounding stator units. Each stator unit contains two MotB subunits that form a central membrane-spanning dimer. The MotB subunits have C-terminal ends that anchor the dimer to the cell wall and N-terminal ends in the cytoplasm. Additionally, each stator consists of five MotA subunits that form a pentamer surrounding the central dimer. The MotA_5_ pentamer has membrane-spanning helices that enclose two proton-conducting channels, one along each subunit of the MotB_2_ dimer.

**Fig. 1.**
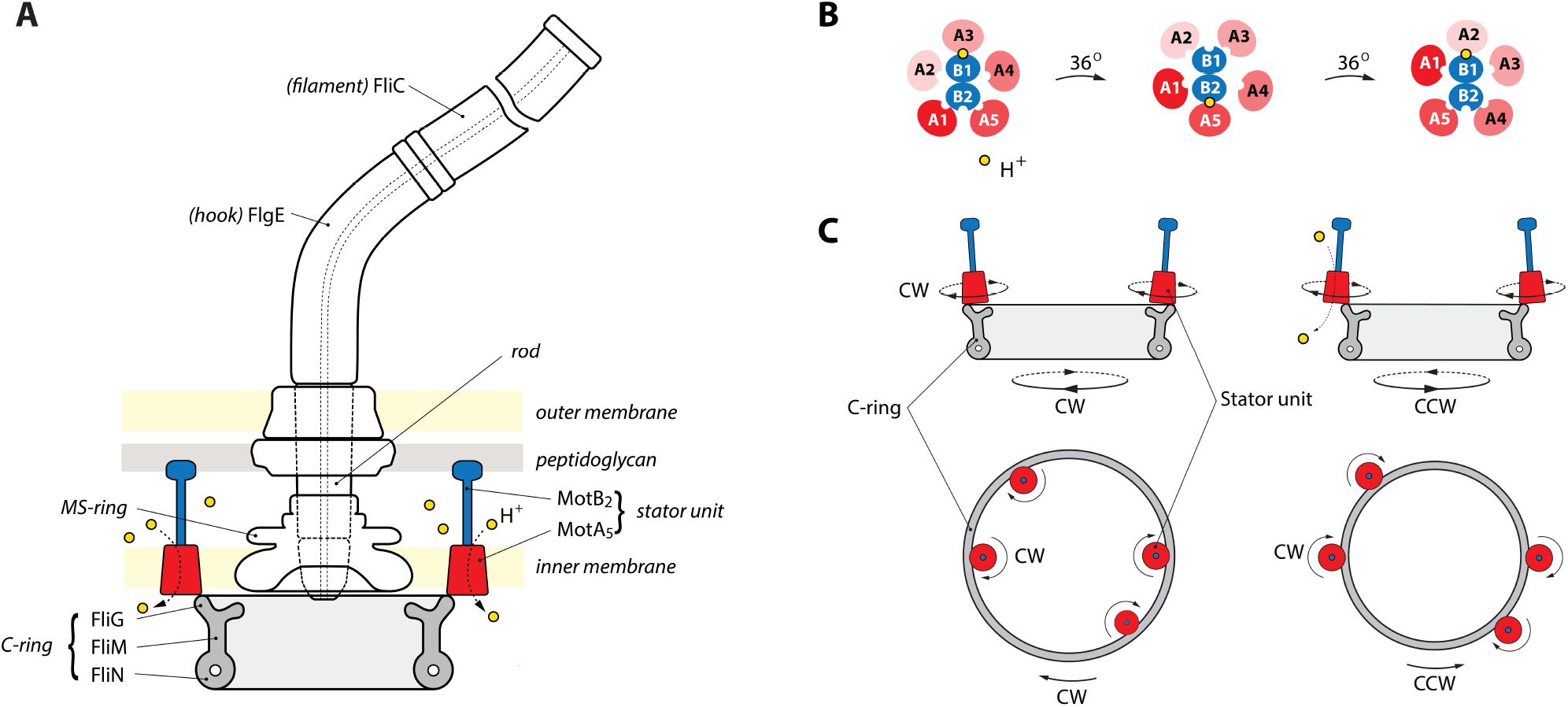
A: Cartoon model of the *E. coli* bacterial flagellar motor. The flagellar filament (made of FliC) is connected via a flexible hook (FlgE) to a central rod with rings in each layer of the cell envelope. The base of the rod is rigidly attached to the C-ring via the MS-ring. The C-ring has 34 subunits of FliG that interact with the MotA subunits of torque-generating stator units. One flagellar motor can accommodate up to 11 stator units.(12) **B: Structural hypothesis for stator rotary movement**, where MotA_5_ rotation around MotB_2_ transmits mechanical power to the C-ring (adapted from (5)). Cartoon of the MotA_5_ /MotB_2_ stator unit viewed from above. One H^+^ channel is open at a time, powering 36°rotation of MotA_5_ around MotB_2_. Two 36°rotations return the two H^+^ channels to their initial open/closed states. **C: Structural hypothesis for bidirectional motor rotation**, driven by unidirectional stator rotation (adapted from (13)). Side and top views of stator units coupled to the rotor (C-ring) in an extended configuration to drive CW-motor rotation (left) and in a compact configuration to drive CCW-motor rotation (right), with CW-stator rotation in both configurations.

Proton-conduction through the stator drives flagellar rotation fueled by an electrochemical gradient across the cell membrane. The structure and stoichiometry of the stator suggests a mechanism where the stator itself might be a rotary motor. (4, 5) Because 5:2 stoichiometry prevents both proton-conducting channels from simultaneously having the same configuration at the MotA_5_/MotB_2_ interface, only one channel can be open at a time. Geometric interplay between five-fold and two-fold symmetric structures predicts that pentamer rotation by 36° around the dimer would toggle the open and closed states of the proton channels. Another 36° rotation in the same direction would reverse the toggle. After two 36° rotations, each MotA subunit will occupy the initial position of its neighboring subunit (Fig. 1B). Thus unidirectional proton-conduction into the cell might directly couple to unidirectional rotation of MotA_5_ pentamer around MotB_2_ dimer.

The mechanical power that drives flagellar motor rotation is transmitted to the C-ring by the cytoplasmic lobes of MotA subunits. The surface of the MotA cytoplasmic lobe has charge complementarity to the surface of the C-ring at specific amino acids on FliG subunits. (7–10) These amino acids represent contact points where the MotA_5_ pentamer can interact with 34 FliG subunits along the C-ring perimeter. If these contact points are where stator and rotor are intermeshed as gearwheels, a conformational change in the C-ring offers a mechanism for bidirectional switching (Fig. 1C). (9–11)

Cryo-EM has revealed different configurations for the C-ring when the flagellar motor is locked in clockwise (CW) or counterclockwise (CCW) modes. In the model of intermeshed gearwheels, unidirectional rotation of MotA_5_ around MotB_2_ might drive bidirectional motor rotation. When the motor is locked in CCW mode, the FliG subunits are poised to contact the innermost MotA subunit of the MotA_5_ pentamer. In this more compact C-ring configuration, when MotA_5_ rotates CW, the C-ring will rotate CCW. When the motor is locked in CW mode, the FliG subunits are poised to contact the outermost subunit of the MotA_5_ pentamer. In this more extended C-ring configuration, when MotA_5_ rotates CW, the C-ring also rotates CW.

Models of torque-generation and bidirectional switching where stator and rotor are intermeshed gearwheels have been based on separate static cryo-EM structures of rotor, stator, and C-ring taken from different bacterial species. (4–6, 9– 11) Here, we sought direct measurement of stator and rotor dynamics in the intact flagellar motor of a living *E. coli* cell.

## Results

### Polarized photo-bleaching microscopy

Fluorescence microscopy using polarized light can probe the spatial orientation of fluorescently-labeled molecules. (14–17) The absorption transition dipole moment (TDM) of a molecular fluorophore makes photon absorption depend on the polarization of excitation light. The probability of photon absorption is proportional to the square of the orthogonal projection of the electric field vector of excitation light onto the TDM direction. The maximum is when the two vectors are parallel and zero when they are perpendicular. The TDM moment also rotates as the molecule rotates. When a rotating fluorophore is excited using light with fixed polarization, fluorescence emission will oscillate twice with each full molecular rotation. Within the rotation plane, the projections of the vector components of TDM and excitation light are twice parallel (at 0° and 180°) and twice orthogonal (at 90° and 270°).

Fusing a fluorophore to every subunit of a radially-symmetric multimer will result in a radial distribution of TDMs. When the fully-labeled multimer rotates around its symmetry axis (*z*), the total fluorescence emission will be nearly constant when excited by light with fixed polarization in the plane of rotation (*x-y*) — when each fluorophore rotates away from peak sensitivity, another fluorophore rotates into place (Fig. 2A, before pulse). However, sensitivity to polarized light can be induced by rapid photo-bleaching using a short, strong pulse of polarized light when the multimer has one angular orientation. A short, strong pulse will mostly bleach the subset of fluorophores with parallel TDMs. After the pulse, fluorescence emission caused by weak excitation with polarized light will oscillate, as unbleached fluorophores rotate in and out of alignment (Fig. 2A, after pulse).

**Fig. 2.**
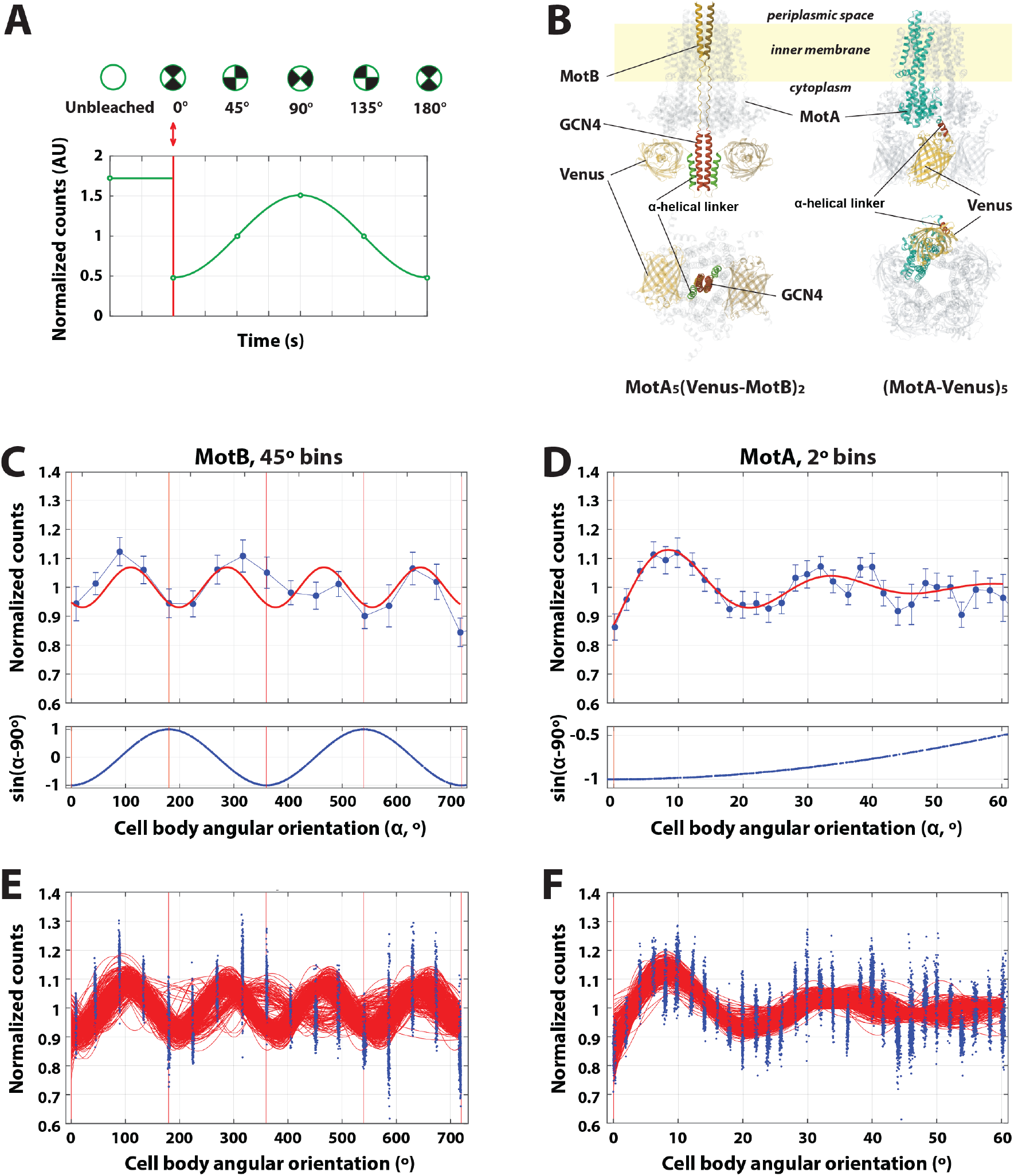
A. Polarized photo-bleaching fluorescence microscopy. Fluorescence emission vs. time (green curve) of a rotating multimer with a radial and symmetric distribution of fluorophore domains. When the unbleached rotating multimer (white circle) is initially probed with weak polarized light, fluorescence emission does not indicate angular orientation. When a strong, short pulse of polarized light is applied (*red double-headed arrow*), a subset of fluorophores is bleached (two black-filled sectors). As the multimer continues to rotate and probed with weak polarized light, the fluorescence emission of unbleached fluophores becomes a reporter of angular orientation. **B. Predicted molecular structures of fluorophore fusions with MotB dimer and MotA pentamer**. (*Left*) AlphaFold predictions of Venus barrel orientations in MotA_5_ (Venus-MotB)_2_ ; only the periplasmic end of MotB is shown). (*Right*) AlphaFold prediction of (MotA-Venus)_5_ One of the MotA-Venus pairs is color coded, the rest are transparent. Side views, along *z*-axis (*top*) and from below, in *x-y* plane of rotation (*bottom*); see also Fig. S7 and Movies 1 and 2. **C**,**E**: **MotB**_**2**_ **experiments**. Immediately after a photobleaching pulse at *t* =0, fluorescence emission dropped and oscillated with twice the frequency of cell body rotation (1.94 *±* 0.18, 45°bins, s.e.m. error bars, *n*=21 motors). **D**,**F: MotA**_**5**_ **experiments**. Immediately after a photobleaching pulse at *t* =0, fluorescence emission dropped and oscillated with periodicity ∼14.6 times faster than cell body rotation (14.59 *±* 1.37, 2°bins, s.e.m. error bars, *n*=36 motors). Top panels (**C, D**) represent mean normalized fluorescence emission (upper) vs. cumulative angular orientation of the tethered cell body (lower), immediately after applying a bleaching pulse of polarized light at angle 0°. Red curved lines represent sinusoidal fits (see **Methods**). Bottom panels (**E, F**) are bootstrap analyses of data in above panels used to calculate confidence intervals (see **Methods**). Each red line is the best fit of single motor traces (blue dots), picked randomly with replacement (*N*=500 iterations).

### Fluorescent labeling of stators in an optimized *E. coli* strain

We made fluorescently-labeled MotA and MotB subunits by fusing Venus (18), an enhanced YFP variant, to each cytoplasmic domain via a linker (Fig. 2B). Tagging stator proteins with large fluorescent proteins might perturb functionality (e.g., rotation direction, speed, switching frequency and CW/CCW bias). These functional parameters might be preserved using longer, flexible or rigid linkers (19). For us, the linker needs to be rigid or short enough to limit the rotational diffusion of Venus with respect to MotA or MotB, but also flexible or long enough to allow large fluorescent tags to position themselves in the crowded environment in a way that preserves functionality. We defined minimal functionality as the ability of the stator to apply torque to the C-ring in a tethered cell.

We used AlphaFold to predict the general structure of the constructs when assembled into stators. We also used AlphaFold to evaluate the orientation of the fluorophores, identify possible points of flexibility and assess their impact on chromophore wobbling when Venus is fused to subunits of either multimer (MotA_5_ or MotB_2_). (20) We then chose the best constructs based on experimental evaluation.

For MotA-Venus, we used a rigid linker (AKEAA)_2_, which extended the *α*-helix at the C-terminus of MotA and formed a continuous helix with the *α*-helix at the N-terminus of Venus. Some flexibility was predicted in MotA, upstream of this *α*-helix. The linkage allows the five Venus monomers to place themselves under the MotA_5_-FliG (Fig. 2B, S7C,F, S8B,D, S10, S11, and **Methods**), clearing the interaction site with the FliG torque coil on the C-ring, assuming the MotA_5_-FliG interaction occurs side-to-side (Fig. S9 and Supporting text MotA-FliG interaction).

For Venus-MotB, we used a highly soluble *α*-helical linker attached to a truncated C-terminus end of Venus (last 6 amino acid residues at the C-terminus end removed). We further extended and constrained the structure by inserting a GCN4 structural motif (21) between the C-terminus end of the *α*-helical linker and the N-terminus end of MotB. In the assembled MotA_5_(Venus-MotB)_2_ stator unit, the two GCN4 helices are predicted to adopt a coiled-coil conformation (via hydrophobic interactions between periodic leucine residues on the opposite GCN4). This is expected to keep the two Venus barrels closer to each other under the MotA_5_ ring, in an antiparallel orientation, preventing their relative rotation about the multiple single covalent bonds in the long N-terminus end of MotB inside MotA_5_ and keeping the interaction site with FliG clear (Figs. 2B, S7A,B,D.E and S13).

We used a background *E. coli* strain deleted for endogenous *motA* and *motB*; *fliC*, the gene for flagellin; and *flgE*, the endogenous gene for hook protein (Fig. 1A). To tether the cells, we designed a “sticky” hook, encoded by a mutant FlgE that adheres spontaneously to clean glass (see **Methods**). When these non-motile cells settle onto a cover slip and self-tether by a single sticky hook, the flagellar motor drives cell body rotation.

To probe rotational motion of MotA_5_, we co-expressed wild-type MotB and the MotA-Venus fusion (strain AP1). To probe MotB_2_, we co-expressed wild-type MotA and the Venus-MotB (strain BP2).

#### Functional evaluation

We qualitatively verified that Venus-MotB restores swimming motility – BP2 cells swim and switch direction when Venus-MotB and MotA are expressed in a Δ*motA*Δ*motB* but otherwise wild type (WT) background strain. BP2 tethers successfully. MotA-Venus does not restore swimming motility when expressed with MotB in a Δ*motA*Δ*motB* background, but AP1 tethers successfully and shows powered rotation, albeit at a lower yield than BP2 or WT. At the same tethered cell density, AP1 provides fewer spinning cells per field of view than BP2, and BP2 fewer than WT. However, when tethered cells spin, both strains reach upper rates comparable to WT. The average speeds of the cells used in these experiments were 2.4 ± 1.5 Hz (mean ± SD, *n*=52) for AP1 or 3.5 ± 2.5 Hz (mean ± SD, *n*=32) for BP2, see also Fig. S16. All tethered cells showed strong CCW rotational bias. Only CCW-rotating motors were analyzed.

Tethered cells typically spin around a bright fluorescent spot, co-localized with the center of rotation observed in bright field, as shown in Fig. S18. The fluorescence emission intensity of the motors spots in the first fluorescence frame is higher in AP1 than in BP2, with an average ratio of (3.3 ± 0.4): (1 ± 0.1) when probed with S-polarized light and (2.8 ± 0.3): (1 ± 0.1) when probed with P-polarized light (mean ± SD; see Table S1).

This ratio between motor fluorescence in AP1 and BP2 roughly agrees with the 2.5:1 (5:2) stoichiometry of MotA:MotB in assembled stators, assuming that each motor is powered by the same average number of stator units in each strain. The emission intensity depends not only on the number of contributing fluorophores in each motor spot, but also on their orientation. The slightly higher than expected MotA-Venus:Venus-MotB emission ratio in S-polarized excitation suggests a higher probability of photo-selection of MotA-Venus (*vs*. Venus-MotB) chromophores when probed in S than in P. This implies a TDM orientation closer to the plane of rotation (*x-y*, probed in S) in MotA-Venus than in Venus-MotB, consistent with our approximation of the TDM orientations based on the AlphaFold predictions (Figs. S7,S8 and Supporting text Approximation of the angle of orientation of the absorption transition dipole moments (TDM)). It is also possible that some Venus tags are cleaved off in the functioning motor, perhaps reducing the overall fluorescence signal without affecting periodicity exhibited by remaining tags.

*E. coli* swimming requires multiple functional motors working in synergy at the same time. Powered tethered rotation only requires one, which seems to successfully assemble in both strains, although at a lower rate in AP1 than in BP2 and, in both, lower then in WT. As both strains exhibit powered tethered rotation around fluorescent motor spots, brighter in AP1 than in BP2 as expected, we conclude that both strains meet the criteria for functional tests of rotational motion.

#### Structural considerations

Structural studies predict that the torque is applied via electrostatic interactions between charged amino acid residues at the cytoplasmic side of the MotA_5_ ring and complementary charged amino acid residues at the cell membrane-facing side of the C-ring. It can be argued that these interactions, between MotA_5_ and FliG_34_ rings (for a 34-fold symmetry C-ring in CCW conformation) occur side-to-side, without overlapping of the two rings along *z*-direction. (See **MotA-FliG interaction site** and Fig. S9 in **Supporting information**).

In all MotA_5_(Venus-MotB)_2_ and (MotA-Venus)_5_ predicted structures, the Venus barrels are located under the MotA_5_ ring (Fig S7). This placement of the fluorescent proteins allows sideways interactions between the complementary charged residues involved in torque coupling distributed along the rims of the MotA_5_ and the FliG_34_ rings (Fig.S12).

In (MotA-Venus)_5_, the Venus barrels themselves do not appear to obstruct the MotA-FliG interface (as they are placed under it), but their partially flexible linkages to MotA might interfere, in some structural predictions. In these instances, such as the prediction depicted in Fig. S10, the Venus barrel would have to slide or move laterally (as in the prediction depicted in Fig. S11) to allow R90 to interact with the D288-D289 pair on FliG. In functional (MotA-Venus)_5_MotB_2_ motors, transient reorientation of Venus barrels at the time of C-ring interaction might lower signal/noise ratio (SNR), without affecting its periodicity. This event would occur in phase with the MotA_5_ rotation, synchronized by the bleaching pulse. Moreover, once a (MotA-Venus)_5_MotB_2_ stator unit is recruited into a motor, the C-ring likely imposes additional steric constraints that further reduce the range of possible fluorophore orientations.

(Venus-MotB)_2_MotA_5_ stators have a lower likelihood of interfering with MotA-FliG interaction site. The Venus-MotB linkage is axial. Also, as they do not rotate with respect to the C-ring (they only translate, tangentially), radially symmetrical orientations around the C-ring that allow unhindered passage of the torque coil and C-ring structures are possible (Fig S13).

### Measuring MotA_5_ and MotB_2_ rotation inside tethered cells

In previous work, we applied photo-bleaching microscopy to unloaded rotating motors lacking flagella. We probed the rotation of fluorescently-labeled FliN subunits (Fig. 1A), and thereby showed that C-ring rotation is locked to the rotation of the extracellular hook. (22) Thus, the C-ring is part of the rotor. Because the C-ring interacts directly with the torque-generating stator unit, the C-ring represents the cytoplasmic gearwheel that rotates the flagellum. (7–10)

Here, we used tethered cells to be able to calibrate motor rotation to stator dynamics. In a tethered cell, the C-ring does not rotate in the laboratory reference frame. As the flagellar motor drives rotation of the tethered cell, the stator units will orbit the C-ring in the lab reference frame. If MotB_2_ is anchored to the cell wall, it should rotate once with each orbit. If MotA_5_ also rotates around MotB_2_, during each orbit MotA_5_ will exhibit additional rotations in the lab reference frame.

In this study, we limited analysis to flagellar motors that rotated exclusively CCW. For the directional-switching model shown in Fig. 1B, MotA_5_ and the tethered cell body are predicted to rotate in the same direction (CW). If MotA_5_ rotation acts on the C-ring like an intermeshed gearwheel, the number of MotA_5_ rotations added to one motor rotation indicates the gear ratio. Thus, in one MotA_5_MotB_2_ stator orbit around the C-ring, the total number of MotA_5_ rotations observed in the lab reference frame would be the gear ratio plus one.

In each experimental trial, we record fluorescence emission from all labeled MotA_5_ or MotB_2_ multimers among several stator units in each flagellar motor. The flagellar motor of a tethered cell typically has up to 10 stator units. (12) Photo-bleaching allows one experimental trial for each tethered cell. To increase signal-to-noise when recording from small numbers of molecules per trial, we need to combine measurements from different tethered cells that typically rotate at different speeds. Our experimental approach is to combine measurements from different cells by calibrating polarization-dependent fluorescence to the angular orientation of each tethered cell. This requires binning, with bin size optimized to provide the highest SNR while retaining appropriate resolution for the expected periodicity.

Each experimental trial begins by delivering a single, strong photo-bleaching pulse of S-polarized light (*<*1.8 msec) to a tethered cell. The electric field vector of S-polarized light is parallel to the rotation plane of the flagellar motor in the tethered cell. The photo-bleaching pulse is followed by a steady train of weak excitation pulses (*<*150 μs) with the same polarization to record fluorescence emission from labeled MotA_5_ or MotB_2_. We simultaneously record the angular orientation of the cell body with bright-field microscopy. This setup allows us to calibrate fluorescence signals from MotA_5_ or MotB_2_ as a function of motor rotation across different cells.

To increase our confidence that the periodic signal is provided by structures labeled with chromophores that have significant TDM orientations in the plane of rotation *(x-y)*, we performed control experiments using P-polarized instead of S-polarized excitation light. In total internal reflection fluorescence (TIRF) mode, the electric field vector of P-polarized light is mostly perpendicular to the microscope slide (see **Methods**). Radially symmetrical structures that rotate around their axis of symmetry do not change their orientation with respect to the axis of rotation and should not produce a periodic signal when probed along *z* (Fig. S7).

To rule out a false positive result related to differences in data analysis (i.e. bin size and angular range), we carried out a complemental analysis. We swapped the MotA and MotB datasets and analyzed each dataset as if belonged to the other group (binning, fitting, initial guess fitting parameters).

#### MotB_2_ rotation is locked to cell body rotation

We tethered cells expressing Venus-MotB and wild-type MotA (strain BP2). To each cell, we applied a single photo-bleaching pulse using S-polarized excitation light. Fluorescence emission, from weak excitation light with the same polarization, dropped immediately and continued to oscillate at twice the angular speed of motor rotation: 1.94 ± 0.18 (mean ± SD, 45° bins, *n*=21; Figs. 2C,E; consistent results using different bin sizes are presented in Fig. S1A-D).

Complemental analysis of MotB data as MotA (smaller bins, for the first 60° of cell body rotation), showed no meaningful significant periodicity (Figs. S4G-L, S6G-I). This range of cell body orientation (0 to 60°) only represents one sixth of a full revolution. Control experiments with P-polarized excitation light showed no consistent periodicity between fluorescence emission and motor rotation (Fig. S2A-F and S3D-F).

We conclude that MotB_2_ rotates around the C-ring once with every motor rotation, consistent with the MotB_2_ dimer being anchored to the cell wall.

#### MotA_5_ rotation is ∼ 6.2x faster than flagellar motor rotation

We tethered cells expressing the MotA-Venus chimera and wild-type MotB (strain AP1). After a photo-bleaching pulse using S-polarized excitation light, polarization-dependent fluorescence emission dropped immediately and continued to oscillate ∼ 14 times faster than motor rotation: 14.44±1.07 (mean ± SD, 2° bins, *n*=36; Fig. 2D,F; similar results for different bin sizes are shown in Fig. S1E-H). In one full 360° cycle of fluorescence emission from (MotA-Venus)_5_, the tethered cell rotates only ∼25°.

Complemental analysis of MotA data as MotB (larger angular bins for a larger range of cell body angular orientations) showed no significant meaningful periodicity (Figs. S4A-F, S6D-F,J-L). As MotA_5_ rotates about twice during each 45° bin, each binned data point is already the average of multiple orientations in all directions. Control experiments using P-polarized excitation light showed no consistent periodicity. (Figs. S2G-L and S3J-L).

We conclude that the MotA_5_ pentamer rotates ∼ 7.2 times per stator orbit around the C-ring of the tethered cell. Subtracting the rotation due to the orbit itself, MotA_5_ rotates ∼6.2 times around MotB_2_ with each motor rotation (6.22 *±* 0.54, mean *±* SD).

## Discussion

We present *in vivo* evidence that rotation of the stator unit in the bacterial flagellar motor — rotation of the MotA_5_ pentamer around the MotB_2_ dimer anchored to the cell wall — is directly coupled to C-ring rotation. Torque-generating stator unit and flagellar motor rotor act like intermeshed gearwheels with ∼ 6.2 gear ratio. Our dynamical observations support structure-based hypotheses that the stator unit itself might be a rotary motor. These hypotheses were based on static cryo-EM structures of rotor, stator, and C-ring taken from different bacteria. (4, 5, 9–11)

Polarized photo-bleaching microscopy with fixed polarization can measure the angular speed but not the direction of MotA_5_ rotation around MotB_2_. Immediately after the photo-bleaching pulse, fluorescence emission peaks when the heteromer turns by 90° in either direction (see Fig. 2A). The direction of stator rotation might be predicted from a structure-based model for bidirectional switching, where CCW rotation of a more compact C-ring is driven by CW rotation of MotA_5_ around MotB_2_ (see Fig. 1C). (9–11)

In the tethered cell, stator and rotor are thought to be strongly-coupled during flagellar motor rotation. (23) If stator and rotor are intermeshed gearwheels, one of the five subunits in each MotA_5_ pentamer should contact one (or an adjacent pair) of the 34 FliG subunits in the C-ring at a time. Each stator work-cycle switches the contact point to the neighboring MotA and FliG subunits. If so, tight-coupling predicts a gear ratio of 34:5 or ∼ 6.8. Gear slippage or skipping would alter the measured gear ratio. Our estimate of ∼ 6.2 for the gear ratio agrees with subunit stoichiometry and nearly tight-coupling.

The oscillating fluorescence signal from MotA_5_ decorrelates more rapidly than the signal from MotB_2_ as the flagellar motor rotates (Fig. 2D, F). In each experimental trial, we record fluorescence signals from all stator units associated with each flagellar motor. We expect all MotB_2_ dimers to be anchored to the cell wall, and so every dimer should rotate at the same frequency in the tethered cell, even if associated with different stators. We expect MotA_5_ pentamers in different stator units to be more likely to rotate at different effective frequencies, as local conditions in different cells (load, proton motive force, number of stators in a motor) might create different gear slippage or skipping conditions. If so, polarization-dependent fluorescence would be most strongly synchronized after the photo-bleaching pulse and gradually decorrelate as stator units rotate out of phase.

The MotA_5_/MotB_2_ stator has structural resemblance to another rotary motor, F1-ATPase, the enzyme that couples proton-conduction across the mitochondrial membrane to ATP synthesis. (24) F1-ATPase drives its catalytic cycle — ADP + P_i_ → ATP — by physically rotating a ring of globular subunits (α_3_β_3_) around a central stalk (γ). Because F1-ATPase is enzymatically active *in vitro*, its rotary mechanism was directly observed by attaching its α_3_ β_3_-subunits to a cover slip and monitoring the rotation of objects attached to the γ-subunit. (25, 26)

We note that the newly reported structures of protein complexes in other bacteria have 5:2 stoichiometry that resembles the MotA_5_/MotB_2_, such as the GldLM complex in *Bacteroidetes* that plays a role in gliding motility and protein secretion and the ExpBD complex in *E. coli* and *Serratia marascens* that plays a role in ligand uptake and active transport. (27) These structures also have molecular ‘axles’ that extend into the periplasm and ‘wheels’ that extend into the cytoplasm. Rotation has not yet been directly observed in other 5:2 protein complexes, but the functional homology of rotational motion observed in MotA_5_/MotB_2_ likely extends to other structurally homologous protein complexes.

Polarized photo-bleaching microscopy provides an *in vivo* probe of molecular dynamics for diverse rotary motors inside the living cell, Here, we demonstrate that the MotA_5_/MotB_2_ torque-generating stator unit constitutes a proton-powered rotary motor that drives the bacterial flagellar motor.

## Materials and Methods

### Polarized photo-bleaching fluorescence microscopy

Our experimental setup (Fig. S15) is modified from earlier setups (22, 28) with improvements described below.

### High-power laser pulses

In this study, we needed to use very short pulses of polarized light (*<*100 μs) to bleach and probe active stator rotation in tethered cells. Short pulses require high power. We implement a 2 W argon ion laser (model no. 2017–06S; Spectra-Physics) tuned to 514 nm and ∼ 650 mW.

Laser pulse duration was controlled with an electro-optical deflector (EOD, model no. 310A; Conoptics). The EOD directed the laser beam through a pinhole (excitation ON) or away (excitation OFF) as a function of input voltage. Input voltage was provided by two high-voltage power supplies (model no. HP 6515A; Hewlett Packard) controlled by a custom-built three-state switch (model no. RIS-688; designed by Winfield Hill, Rowland Institute at Harvard).

Laser pulse amplitude was controlled with an electro-optical modulator (EOM; model no. 350–50; Conoptics) driven by a high-voltage differential amplifier (model no. 302 RM; Conoptics) followed by a Glan-laser polarizer (part no. GL10-A; Thorlabs) oriented in S. The EOM changes the polarization of output light as a function of applied voltage, from P to S to different degrees of elliptical to circular polarization. The EOM delivered maximum attenuation between excitation pulses and maximum intensity during the bleaching pulse. For experiments with excitation and bleaching in P, a half-waveplate (part no. WPH05M-514, Thorlabs) was introduced in the optical path after the Glan polarizer, followed by a second Glan polarizer oriented in P. For experiments on beads, with excitation in different polarizations during the same sequence acquisition, all Glan polarizers were removed from the setup and the EOM was fed with the appropriate S, P or C (for circularly polarized output) voltage during each camera exposure.

Laser light was diverted into the sample with a long-pass dichroic mirror (part no. Di01-R514-25x36; Semrock). A 514 nm single-notch filter (part no. NF01-514U-25; Semrock) on the emission side of the optical path was used to attenuate scattered excitation light. Fluorescence emission was diverted to an EMCCD camera (Andor DU-860E; Andor Technology).

A microcontroller generated time and amplitude control pulses for fluorescence excitation and bright field illumination. The microcontroller controlled the EMCCD camera using Andor Solis software and phase camera using Labview (NI). Synchrony with EMCCD camera exposure was monitored via its “Fire” output. The EMCCD camera was set to acquire alternately bright field and fluorescence exposures. (29)

### Light microscopy

We added capacity for bright-field and phase-microscopy to allow simultaneous cell tracking. The light source for cell tracking was a red/amber (613nm) Luminus Phlatlight PT-54 RAX LED driven by a custom built driver. A beam splitter with a short pass (∼ 600 nm) dichroic mirror mounted in the emission path diverted LED light towards the phase camera (DCC1240M CMOS camera; Thorlabs) for focusing, orientation on the slide and cell tracking.

### Imaging tethered cells

Tethered cells in tunnel slides were imaged with either a CFI60 Plan Fluor 40x 1.3 N.A. oil immersion objective (in epi-fluorescence) or a CFI60 Apochromat 60x 1.49 N.A oil immersion objective (in total internal reflection fluorescence, TIRF) on a Nikon TE300 inverted microscope, modified to allow laser excitation from the side.

## Bacterial strains, cell cultures and sample preparation

### Molecular biology

The strains used in this study are derived from the motile wild type (WT) strain MG1655. (30) Deletions of *motB, motA, fliC*, and *flgE* were made sequentially, alone, and in combination (Δ*motA*Δ*motB*) using the Datsenko and Wanner method with pKD3/pKD4 plasmids. (31) An 85 bp scar remained after Flp/FRT recombination to eliminate the antibiotic resistance genes. Deletions were verified using PCR and Sanger sequencing of the PCR products.

We expressed a modified FlgE variant that spontaneously adheres to glass from a plasmid with kanamycin resistance, in both BP2 and AP1 strains. We based the design of our “sticky hook” on a previous insertion of the AviTag protein sequence into the flagellar hook protein FlgE (*FlgE C Avi*). (32, 33) Specifically, we modified the AviTag sequence to include sb7p (RQS**Φ**RGR), a less hydrophylic variant of the original silica binding peptide 7 (S replaced with F in position 4). (34)

We made the BP2 strain for MotB experiments by transforming Δ*motA*Δ*motB*Δ*f liC*Δ*f lgE* cells with a pTrcE plasmid (with ampicillin resistance) carrying both WT *motA* and *Venus*-(L9:48-64)-GSGGS-*GCN4-motB* construct. L9:48-64 is a a highly soluble *α*-helical linker (EAQKQKEQRQAAEELAN), identified in (35). GCN4 is a “leucine zipper”, intended to promote dimerization with the GCN4 motif on the other Venus-MotB monomer of the MotB2 stalk. The first 6 amino acid residues at the N-terminus end of MotB were removed.

We made the AP1 strain for MotA experiments by transforming Δ*motA*Δ*motB*Δ*f liC*Δ*f lgE* cells with a pTrcE plasmid carrying both WT *motB* and *motA*-(AKEAA)_2_-*Venus* (see Table S1 for the complete sequences).

### Cell culture

TB cell cultures started from a saturated LB cell suspension stock (1:20) supplemented with antibiotics. Cells were grown in a shaking incubator at 30° C for about 4 h to 0.5-0.6 optical density at 600 nm. An 0.5 ml aliquot was washed three times by centrifugation and resuspension in 1ml motility buffer (10mM potassium phosphate, 0.1mM EDTA, 10mM lactate, pH 7.0).

About 30 μl of cell suspension was loaded in “tunnel” slides, prepared from coverslips (No. 1.5, VWR Scientific) separated by two layers of double-sided Scotch tape. The loaded tunnel slides were placed in a humidity chamber at room temperature for ∼ 15min, to allow cells to sediment and self-tether. Untethered cells were washed out with motility buffer. Tethered cells were given 30-45 min to allow maximum stator recruitment to the flagellar motor. (36–38)

### Experiments on fluorescent beads

We confirmed that our setup is able to selectively bleach or image with S-or P-polarized light by testing with commercially-available fluorescent beads adsorbed to tunnel slides. About 40 μl of poly-L-lysine (0.01%; Sigma-Aldrich) were loaded in the tunnel slide, incubated for ∼ 5min, and washed out with water. Fluorescent 1 μm beads suspended in water were loaded into the slide, and allowed to settle to the poly-L-lysine treated glass for ∼ 15 min. Excess beads were washed out. This sample was probed alternately with S- and P-polarized light synchronized to camera exposure (all odd frames were excited in P, all even frames were excited in S), bleached in S, and probed again with alternating S (even frames) and P (odd frames) polarization (Fig. S17A,E). For bleaching in P, a half-waveplate was inserted in the optical path, after the EOM, reversing the polarization of each frame (all even frames were excited in P, al odd frames were excited in S) and of the bleaching pulse (Fig. S17B,F). These experiments confirmed that our setup delivers photo-bleaching and excitation pulses with selective polarization (either S or P) with both high N/A objectives. Probing with circularly polarized light (C), which does not excite fluorophores preferentially by their orientation, resulted in similar emission decaying after bleaching in either S or P (Fig. S17C,D,G,H).

### Experiments on cells

To probe polarization in the rotational plane of the tethered cell, we used S-polarized photo-bleaching and excitation pulses (i.e., laser light with electric-fields parallel to the *x-y* plane of motor rotation). In both epi-fluorescence and TIRF microscopy, S-polarization of incident light is preserved from the input aperture of the objective to the sample (slide).

In control experiments, we provided P-polarized light (with electric-fields parallel to the axis of motor rotation) using TIRF mode. The evanescent wave of TIRF microscopy is elliptically polarized in the *x-z* plane of incidence, with the stronger component in *z* direction (perpendicular to the slide and parallel to the axis of rotation) and weaker component (∼ 14-fold less intense) in the *x* direction. As the rotating fluorophores do not change orientation with respect to the *z*-axis of rotation and most excitation is provided by the *z*-component of the evanescent wave, fluorescence emission in these control experiments should not show significant periodicity with respect to motor rotation.

The experimental parameters were optimized empirically for each strain. For each experiment, a total of 2500 combined fluorescence/bright field frames were acquired at rates of 505.21 frames/s (128×128 pixels) or 909.09 frames/s (128×64 pixels) in “frame transfer” mode, which provided an almost continuous camera acquisition with ∼ 1.8 ms or ∼ 1.08 ms exposures per frame and ∼ 20 μs pauses between frames. Each frame was designated as either “fluorescence” or “bright field” prior to the acquisition, with a constant ratio fluorescence:bright field of 1:4, 1:6 or 1:10 for the entire acquisition. For probing, a laser excitation pulse of either 50 or 150 μs was applied 50 μs after the start of each fluorescence frame. For polarized bleaching, a laser pulse of either 800 μs or 1800 μs of the same polarization was applied during frame 45 or 95. A red light illumination pulse of 300 μs was triggered 50 μs after the start of each bright field frame. A combined fluorescence/bright field acquisition of a tethered cell rotating around a fluorescent spot is depicted in Fig. S18 (see also Movie 4).

### Data analysis

For each experimental trial, fluorescence emission and motor orientation were quantified. Bright field images immediately before and after each fluorescence image were averaged to determine the angular orientation of the cell body synchronized to the fluorescence image. Cell body orientation was determined by orthogonal linear regression to the image of the cell body. The fitted slope was converted to angular orientation. The center of rotation was determined either from the center-of-mass or from the center of a two-dimensional Gaussian fit to an image created by averaging together all bright field frames. Alternatively, the cell body orientation was calculated by fitting a circle to the points corresponding to the center of mass of the cell body in each frame, plotted for all the frames. The angular orientation in each frame was calculated as the standard position angle defined by the point on the fit circle that corresponds to the center of mass of the cell body. Both methods provided similar results. The flagellar motors of all tethered cells had strong CCW bias. Only cells that rotated exclusively CCW were studied. The motors that switched the direction of rotation during the relevant part of the acquisition (the first 60° of cell body rotation for MotA experiments or the first 730° of cell body rotation for MotB experiments were discarded from analysis).

Fluorescence emission from each flagellar motor was quantified as the sum of all pixel counts within a radius of a 150 nm mask around the point representing the location of the motor, after subtracting the background value of each pixel. Flagellar motor location was determined by averaging all fluorescence frames and fitting a 2-dimensional Gaussian to a cropped area centered on tethered motor using bright field images. Background pixel values (mostly dark noise in the EMCCD) were obtained by averaging sequences of 10,000 frames of tunnel slides with motility buffer but no cells, acquired after each experiment under identical conditions, using the same camera acquisition and laser excitation parameters.

Fluorescence emission vs. motor angular orientation data sets across experimental trials were combined. The fluorescence emission vs. time curves were fit and normalized to a single exponential decay with offset, *y* = *ae*^−*bx*^ + *c*, to compensate for the bleaching. Normalized fluorescence emission vs. angular orientation datasets were created. The angular orientation for each frame in a sequence was adjusted by subtracting the orientation at the time of bleaching in that sequence. Thus, bleaching always occurred at 0° in every trial.

Absolute angular orientation in each frame (from 0° - 360°), was converted to cumulative angular distance traveled. After binning in intervals of angular distance, average normalized fluorescence emission as a function of average cumulative angular orientation was calculated and plotted. Each curve was fit with sinusoidal oscillations with free parameters for amplitude (*a*), angular frequency (*c*), phase shift (*d*), and exponential decay (*b*):

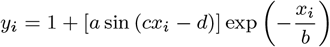

where *y*_*i*_ is the amplitude of the fluorescence signal in bin *i, a* is the initial amplitude of the periodic signal, *x*_*i*_ is the angular orientation of the cell body in bin *i, c* is the multiplication factor of the rotation rate of the structure under investigation with respect to the rotation rate of the cell body. Hence, *cx*_*i*_ corresponds to the angular orientation of the structure under investigation. *d* is the phase shift with respect to the bleaching angle, and is expected to be *π*/2. *b* is a decay constant to account for fluorophore bleaching or signal decorrelation.

The quality of fit was evaluated and confidence intervals were assessed using bootstrap methods. (22, 39) A number of *N* synthetic datasets composed of *n* normalized fluorescence emission vs. cumulative cell body angular orientation curves were constructed by randomly drawing *n* datasets from the original *n* datasets in each experimental group with replacement. Each synthetic dataset was processed in the same way as the original dataset (i.e., binning and fit), resulting in *N* fitting curves and *N* sets of fitting parameters. We chose *N* =500.

Confidence intervals were calculated as the percentage of synthetic datasets that provided fitting parameters within a range of values around the fit of the original dataset (see Fig. S3). These values were a frequency multiplier parameter between 1x and 3x (for MotB_2_ experiments) or between 10x and 20x (for MotA_5_ experiments), a phase shift between 25° and 145° and a normalized initial amplitude between 0.05 and 0.5 (for both MotB_2_ and MotA_5_ experiments). Confidence intervals were used to calculate standard deviation and mean. Data analysis was done with custom scripts in Matlab (Mathworks, Inc.) and Labview (NI).

## ACKNOWLEDGMENTS

This work was supported by NSF-2146519 and the Dean’s Competitive Fund at Harvard University. We are further grateful for the support of Deans Christopher Stubbs and Jeff Lichtman.

