## Supporting Information for "Torque-generating units of the bacterial flagellar motor are rotary motors"

#### This PDF file includes:

Supporting text
Figs. S1 to S18
Tables S1 to S2
Legends for Movies S1 to S4
SI References

#### Other supporting materials for this manuscript include the following:

Movies S1 to S4

### Supporting Information Text

### MotA-FliG interaction

Structural studies predict that the torque is applied via electrostatic interactions between charged residues at the base of $MotA_5$ and complementary charged residues in the FliG "torque coil", at the upper side (cell-membrane facing) of the C-ring.

In *E. coli*, pairs of positively charged R90 and negatively charged E98 residues at outer rim of the cytoplasmic side of MotA interact with pairs of positively charged R281 and negatively charged D288-D289 residues in the FliG torque coil.

Alternating positively and negatively charged residues along the rims of $FliG_{34}$ and $MotA_5$ rings are not uniformly distributed. Within each FliG torque coil, the complementary charges (R281 and D288-D289) are spaced at ∼1 nm. In a C-ring with a 34-fold symmetry locked in CCW conformation, adjacent FliG torque coils are spaced at ∼4 nm, the same pitch as the negatively charged residues (E98) in $MotA_5$, with the positively charged R281 further away from the center of rotation of the C-ring. In each MotA, the complementary charges (R90 and E98) are spaced at ∼2 nm, with the negatively charged E98 further away from the center of rotation.

At any location of FliG along the CCW $FliG_{34}$ ring, the positively charged R281 is slightly "ahead" (in CW direction) of its closest D288-D289 negatively charged neighbor, as viewed from outside the cell. In contrast, in a $MotA_5$ ring viewed from the same perspective, any negatively charged E98 residue is slightly ahead (in CW direction) of its closest positively charged neighbor. Thus, electrostatic interactions between $MotA_5$ and $FliG_{34}$ resemble intermeshed gearwheels (a cogwheel model), in which each charged residue of a cog in a ring interacts with complementary charges on different, adjacent cogs on the other ring (see Fig. S9B).

When observed from the side, all complementary charged residues in a ring are located in different, parallel $x$-$y$ planes along the $z$-axis of rotation. In $MotA_5$, all negatively charged E89 residues are located in a plane closer to the cell membrane than the plane of positively charged R90 residues. In $FliG_{34}$, all positively charged R281 residues are located in an $x$-$y$ plane closer to the cell membrane than the plane of negatively charged residues (see Fig. S9D)

In conjunction with the cogwheel model, this implies that interactions between $MotA_5$ and $FliG_{34}$ occur **side-to-side** (as opposed to up-to-down, a conformation in which $MotA_5$ would apply torque to the C-ring from above). Thus, labeling stator units with fluorescent proteins located under the $MotA_5$-ring allows the construction of functional mutants that do not interfere with torque transmission at the MotA-FliG interface.

### Approximation of the angle of orientation of the absorption transition dipole moments (TDM) in MotA-Venus and Venus-MotB constructs

We used AI-based structural predictions to better understand the relative TDM orientations of each Venus tag with respect the axis of rotation ($z$), the $x$-$y$ plane of rotation, and the other Venus tags in each construct.

We approximated the orientation of the absorption transition dipole moments (TDM) by an imaginary line that connects the amino acid residues Val120 and Ser147 on opposite sides of the Venus barrel (Fig. S14). We found (by examining the published crystal structure of Venus (1) using the web-based viewer Mol (Molstar):*) that this line is reasonably parallel to the line that connect the two farthest C atoms in the chromophore aromatic rings (rigidly attached to the central alpha-helix), reported as a heuristic rule to predict the orientation of TDM in fluorescent proteins (2).

Although this line is not expected to be an accurate prediction of TDM, it can be easily added to each construct representation. This allows comparing Venus barrel orientations across different constructs, or across different predictions of the same construct. The latter can be used to estimate the degree of wobbling in a given construct (see Fig.S8). The Venus barrels in a given construct can change their orientation with respect to the tagged MotB or MotA by translating or rotating about a flexible region in their linkage, within a range of angles limited by local sterical constraints. Given the much shorter fluorescence emission lifetime of Venus (3 ns) during a single fluorescence exposure (50-150 µs) the chromophore is expected to visit all angles within the range, some with a higher probability than others, lowering the fluorescence anisotropy and the signal to noise ratio (SNR).

To calculate the angular orientations of the chromophore, we used its surrogate projection on the three $x$-$y$, $x$-$z$ and $y$-$z$ planes when the $(MotA\text{-}Venus)_5$ or $(Venus\text{-}MotB)_2$ structure was oriented with the expected axis of rotation parallel to $z$-axis.

The range of angles estimated for the predictions in Fig.S8 in the $x$-$y$ plane of rotation (probed with S-polarized light, along $x$ axis) is 46.6° (0° ± 23.3°) for $(MotB\text{-}Venus)_2$ and 28.9° (0° ± 14.5°) for $(MotA\text{-}Venus)_5$. The range of angles along $z$-axis of rotation (probed with P-polarized light) is 45.9° to 71.5° for $(MotB\text{-}Venus)_2$ and 71.9° to 95.9° for $(MotA\text{-}Venus)_5$. The probability of exciting the fluorophores with P-polarized light is higher in $(Venus\text{-}MotB)_2$ (∼0.3 to ∼0.7) than in $(MotA\text{-}Venus)_5$ (∼0.05 to ∼0.3). In the $x$-$y$ plane of rotation, the probability of exciting chromophores with S-polarized light is higher for $(MotA\text{-}Venus)_5$ (∼0.9 to ∼1) than for $(Venus\text{-}MotB)_2$ (∼0.7 to ∼0.9). These approximations are consistent with the fluorescence emission intensity data in Table S1.

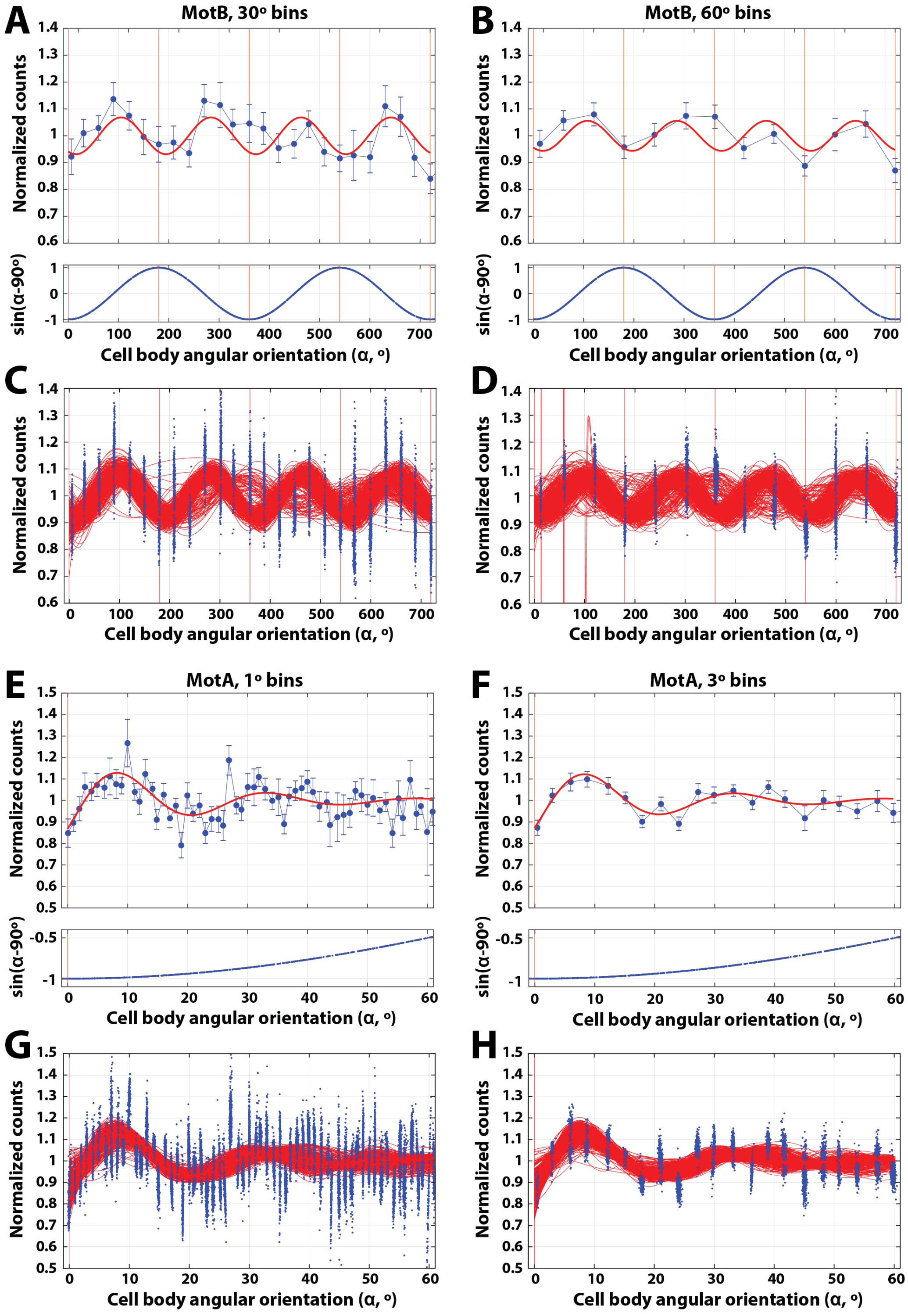


**Fig. S1. A-D: MotB results (30° and 60° binning).** After a photo-bleaching pulse at $t$=0, subsequent polarization-dependent fluorescence emission oscillated at twice the angular speed of motor rotation. When data is averaged in 30° bins (**A**,**C**), we estimate a multiplication factor of 1.95 ± 0.19. When data is averaged in 60° bins (**B**,**D**), we estimate a multiplication factor of 1.95 ± 0.19 (mean ± SD), $n$=21 motors. **E-F: MotA results (1° and 3° binning).** After a photo-bleaching pulse at $t$=0, subsequent polarization-dependent fluorescence emission oscillated with periodicity about 14.9 times faster than cell body rotation when data is averaged in 1° bins (14.87 ± 1.32, mean ± SD) or 14.6 when data is averaged in 3° bins (14.59 ± 1.23, mean ± SD), $n$=36 motors. In both experiment groups (MotB and MotA), top panels (**A**,**B**,**E**,**F**) represent mean normalized fluorescence emission vs. cumulative angular orientation of the tethered cell body, immediately after applying a bleaching pulse of polarized light at angle 0° (upper), and phase-shift adjusted sine curve of the cell body orientation vs. its cumulative angular orientation (lower). Red curved lines represent sinusoidal fits (see **Methods**). Bottom panels (**C**,**D**,**G**,**H**) are bootstrap analyses of data in above panels used to calculate confidence intervals (CI). Each red line is the best fit of single motor traces (blue dots), picked randomly with replacement ($N$=500 iterations).

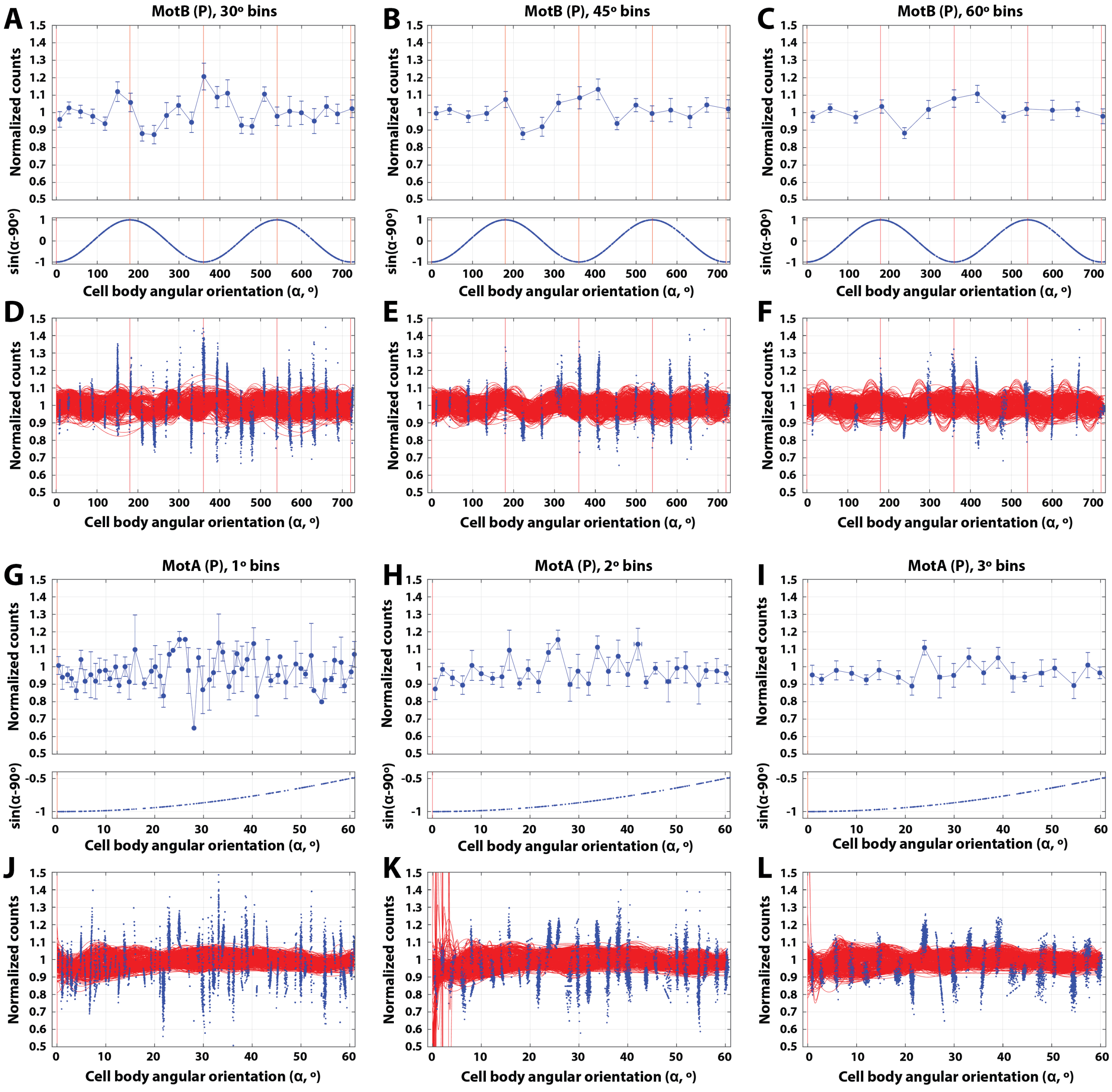


**Fig. S2. A-F: MotB control results.** No consistent periodicity between fluorescence emission and motor rotation is observed for excitation along $z$-axis of rotation (see **Methods**) in datasets binned in 30° (**A**,**D**), 45° (**B**,**E**) and 60° bins (**C**,**F**); $n$=11 motors. **G-L**: **MotA control results.** No consistent periodicity between fluorescence emission and motor rotation is observed for excitation along $z$-axis of rotation in datasets binned in 1° (**G**,**J**), 2° (**H**,**K**) and 3° bins (**I**,**L**); $n$=25 motors. Panels **D-F** and **J-L** show bootstrap analyses of data in the panel above (**A-C** and **G-I**, respectively) used to calculate CI. Each red line is the best fit of single motor traces (blue dots), picked randomly with replacement ($N$=500 iterations).

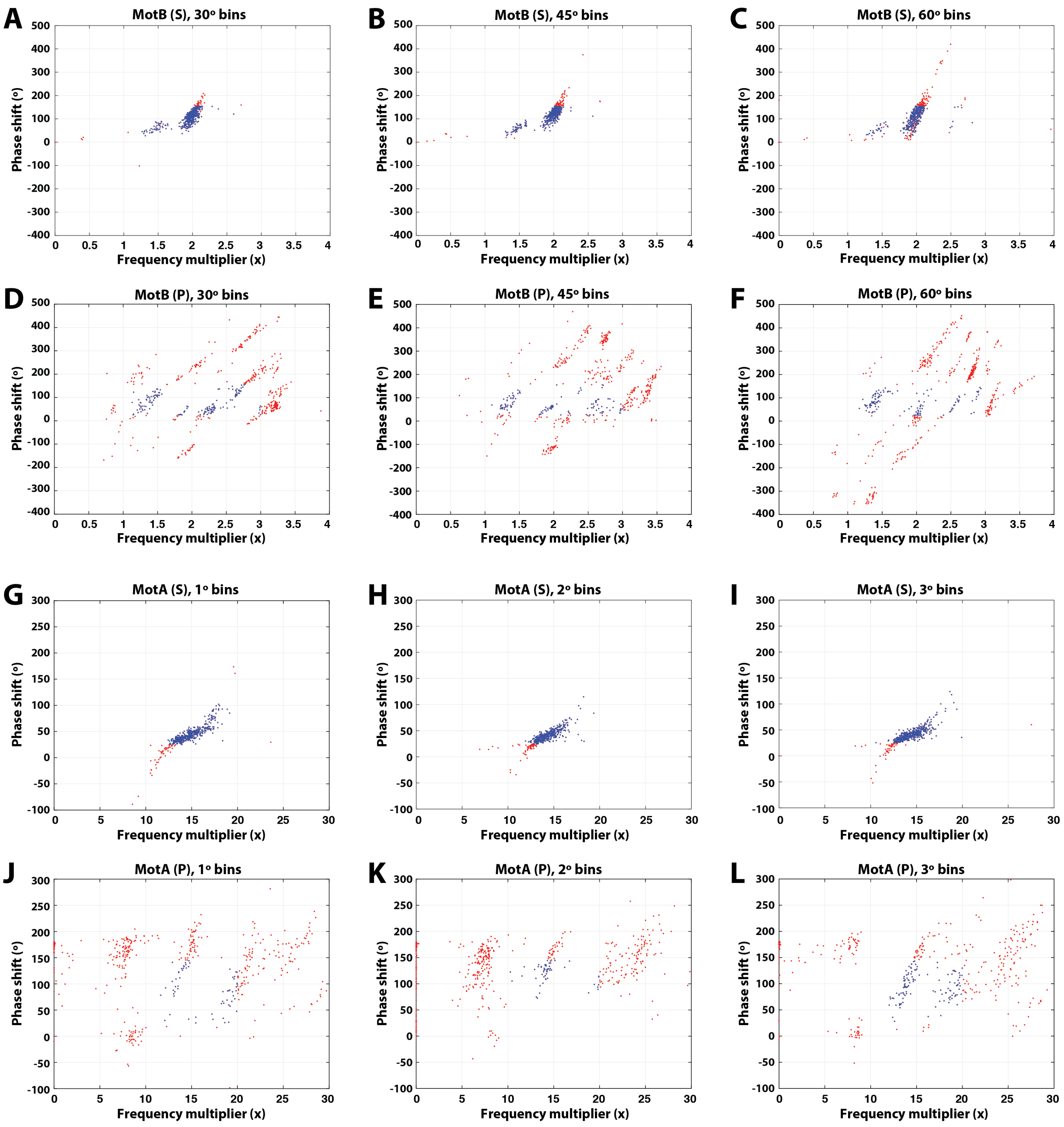


**Fig. S3. MotB (A-F) and Mot A (G-L) confidence intervals.** Experiments with fluorescence excitation using S-polarized (**A**-**C**, **G**-**I**) or P-polarized (**D**-**F**, **J**-**L**) light, for datasets binned in 30° (**A**,**D**), 45° (**B**,**E**) and 60° bins (**C**,**F**) for MotB and 1° (**G**,**J**), 2° (**H**,**K**) and 3° bins (**I**,**L**) for MotA. Best fit parameters (phase shift and frequency multiplier) for the fit lines in Figs. 2E,F, S2D-F, andS2J-L, that jointly fall within (blue dots) or outside (red dots) limits of a positive result (frequency multiplier, appropriate phase shift, significant amplitude - see **Methods**). **A**: CI=91.4%; **B**: CI=85.8%; **C**: CI=70.4%; **D**: CI=29.8%; **E**: CI=23.8%; **F**: CI=26.4%; **G**: CI=90.4%; **H**: CI=88.4%; **I**: CI=89.8%; **J**: CI=12.2%; **K**: CI=11.0%; **L**: CI=24.0%.

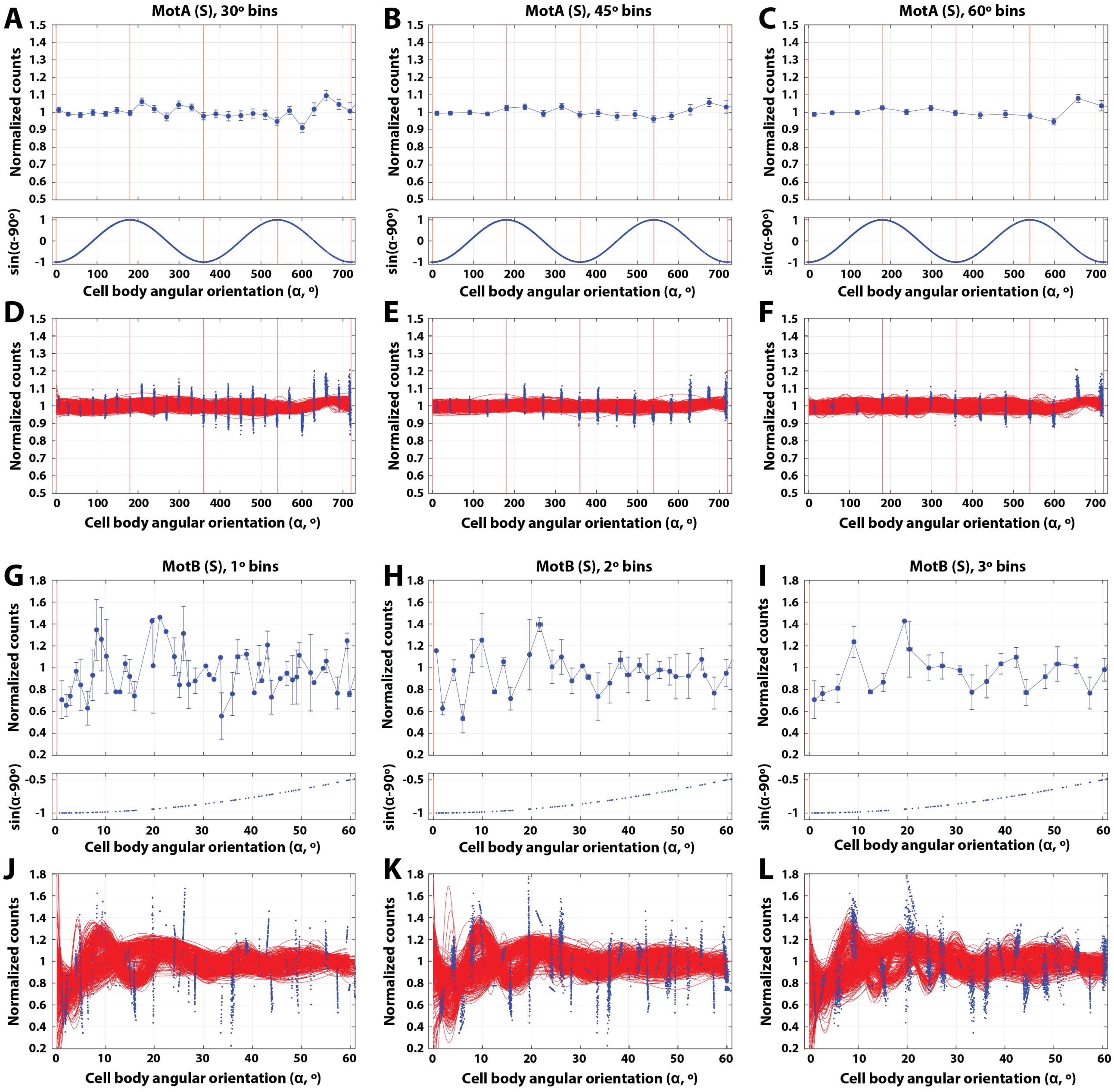


**Fig. S4. Complemental analysis, excitation with S-polarized light**. **A**-**F**. **MotA data analyzed as MotB**. No consistent/meaningful periodicity between fluorescence emission and motor rotation is observed if the Mot B dataset is binned in 1° (**A**,**D**), 2° (**B**,**E**) and 3° bins (**C**,**F**); *n*=36 motors. **G**-**L**: **MotB data analyzed as MotA**. No consistent/meaningful periodicity between fluorescence emission and motor rotation is observed if the MotA dataset is binned in 45° (**G**,**J**), 30° (**H**,**K**) and 60° bins (**I**,**L**); *n*=21 motors. Panels **D**-**F** and **J**-**L** show bootstrap analyses of data in the panel above (**A**-**C** and **G**-**I**, respectively) used to calculate CI. Each red line is the best fit of single motor traces (blue dots), picked randomly with replacement (*N*=500 iterations).

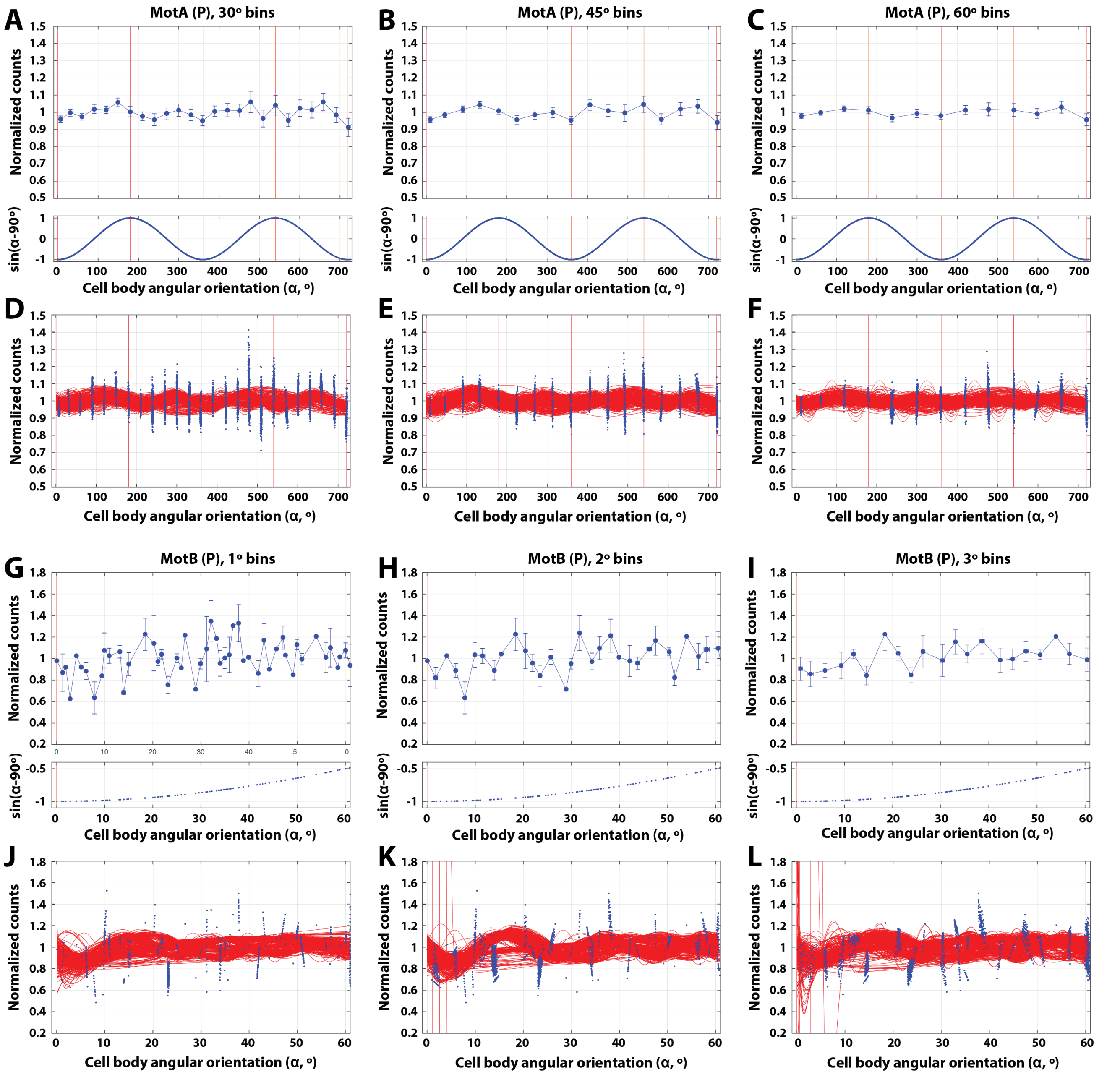


**Fig. S5. Complemental analysis, excitation with P-polarized light (control)**. **A**-**F**: **MotA data analyzed as MotB**. No consistent/meaningful periodicity between fluorescence emission and motor rotation is observed if the MotA dataset is binned in 45° (**A**,**D**), 30° (**B**,**E**) and 60° bins (**C**,**F**); *n*=16 motors. **G**-**L**: **MotB data analyzed as MotA**. No consistent/meaningful periodicity between fluorescence emission and motor rotation is observed if the MotA dataset is binned in 1° (**G**,**J**), 2° (**H**,**K**) and 3° bins (**I**,**L**); *n*=11 motors. Panels **D**-**F** and **J**-**L** show bootstrap analyses of data in the panel above (**A**-**C** and **G**-**I**, respectively) used to calculate CI. Each red line is the best fit of single motor traces (blue dots), picked randomly with replacement (*N*=500 iterations).

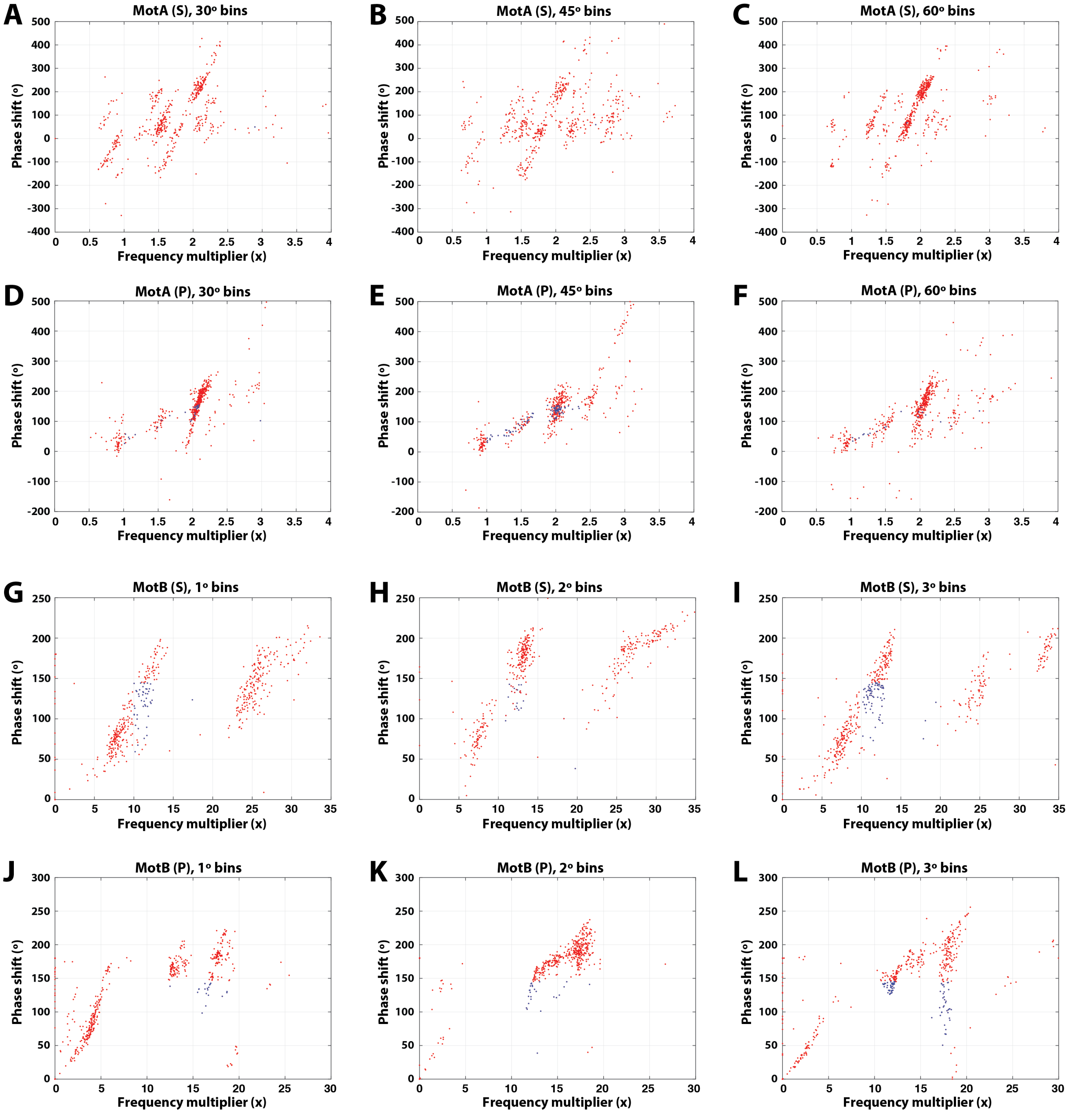


**Fig. S6. Complementary analysis: confidence intervals.** **A**-**F**: MotA data analyzed as MotB, with fluorescence excitation with S-polarized (**A**-**C**) or P-polarized (**D**-**F**) light. **G**-**L**: MotB data analyzed as MotA, with fluorescence excitation with S-polarized light (**G**-**I**) or P-polarized light (**J**-**L**). Datasets binned in 1° (**A**,**D**), 2° (**B**,**E**) and 3° bins (**C**,**F**) for MotB and 30° (**G**,**J**), 45° (**H**,**K**) and 60° bins (**I**,**L**) for MotA. Phase shift and frequency multiplier best fit parameters, for data depicted in Figs. S15D-F,J-L and S5D-F,J-L, that jointly fall within (blue dots) or outside (red dots) limits corresponding to a positive result (frequency multiplier, appropriate phase shift, significant amplitude - see **Methods**). **A**: CI=7.6%; **B**: CI=3.2%; **C**: CI=13.4%; **D**: CI=14.8%; **E**: CI=31.8%; **F**: CI=2.8%; **G**: CI=2.6%; **H**: CI=8%; **I**: CI=5.8%; **J**: CI=14.8%; **K**: CI=31.8%; **L**: CI=28.0%.

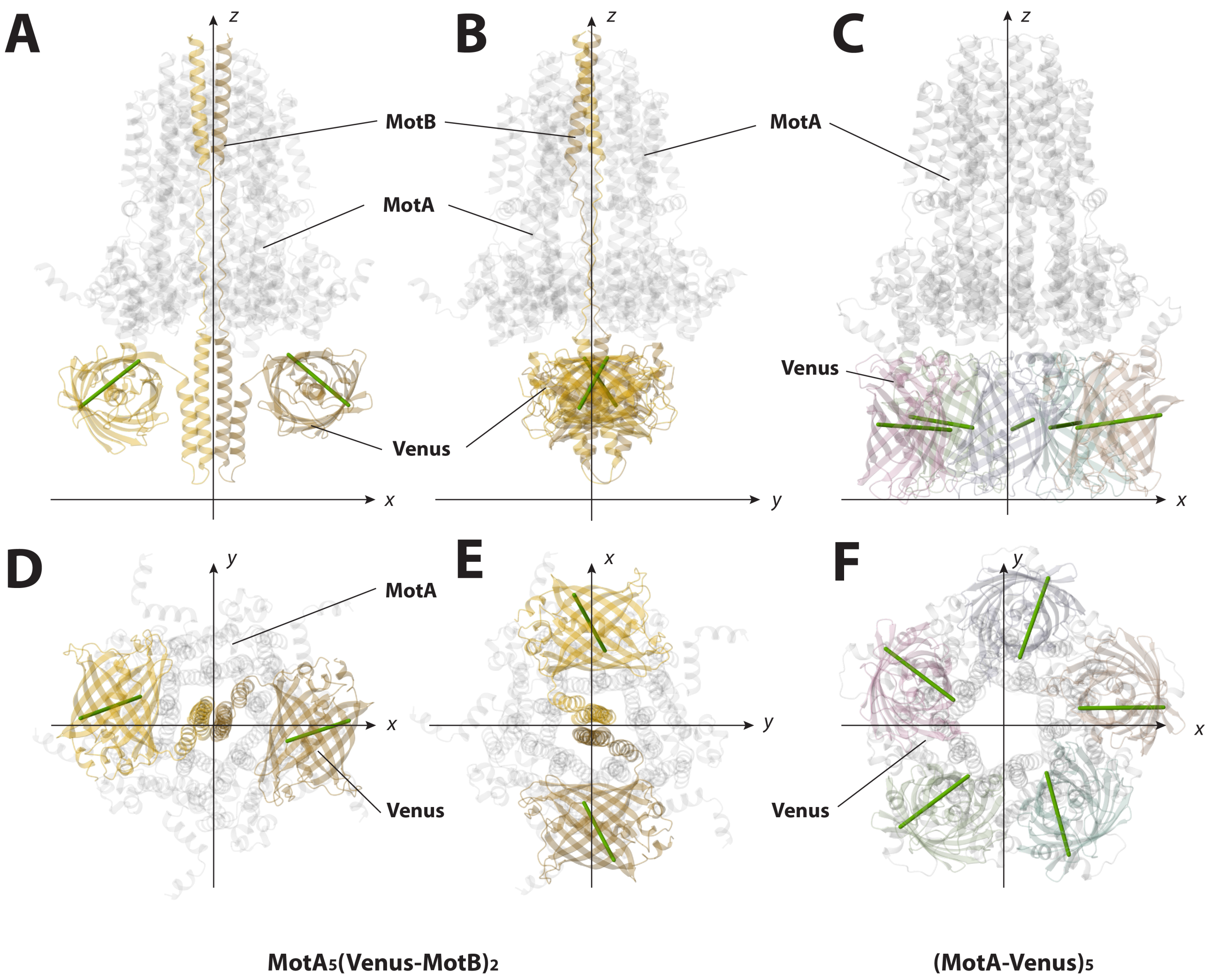


**Fig. S7. Radial orientation of chromophores**. Approximations of the chromophore orientations (thick green lines across the Venus barrels) in cartoon representations of predicted structures. *Top panels:* Side views of $MotA_5$(Venus-MotB)$_2$ (**A**), rotated with 90°along the vertical ($z$) axis (**B**) and of (MotA-Venus)$_5$ (**C**). *Bottom panels:* structures in the panels above, viewed from the bottom ($x$-$y$ plane). The probability of exciting a chromophore with polarized light with electric vectors parallel with a given axis is proportional with the square orthogonal projection of the absorption transition dipole moment (TDM) on that axis. According to the orientations depicted in this figure, the chromophores have a significant projection on the $x$-$y$ plane of rotation in both constructs. In (MotA-Venus)$_5$, they are radially oriented. In $MotA_5$(Venus-MotB)$_2$, they show a parallel orientation in $x$-$y$ plane, inferring that polarized photo-bleaching is neither required nor efficient to probe rotation in single stator units. However, multiple stator units powering a C-ring, probed simultaneously would have have multiple chromophores with radial distributions in both constructs.

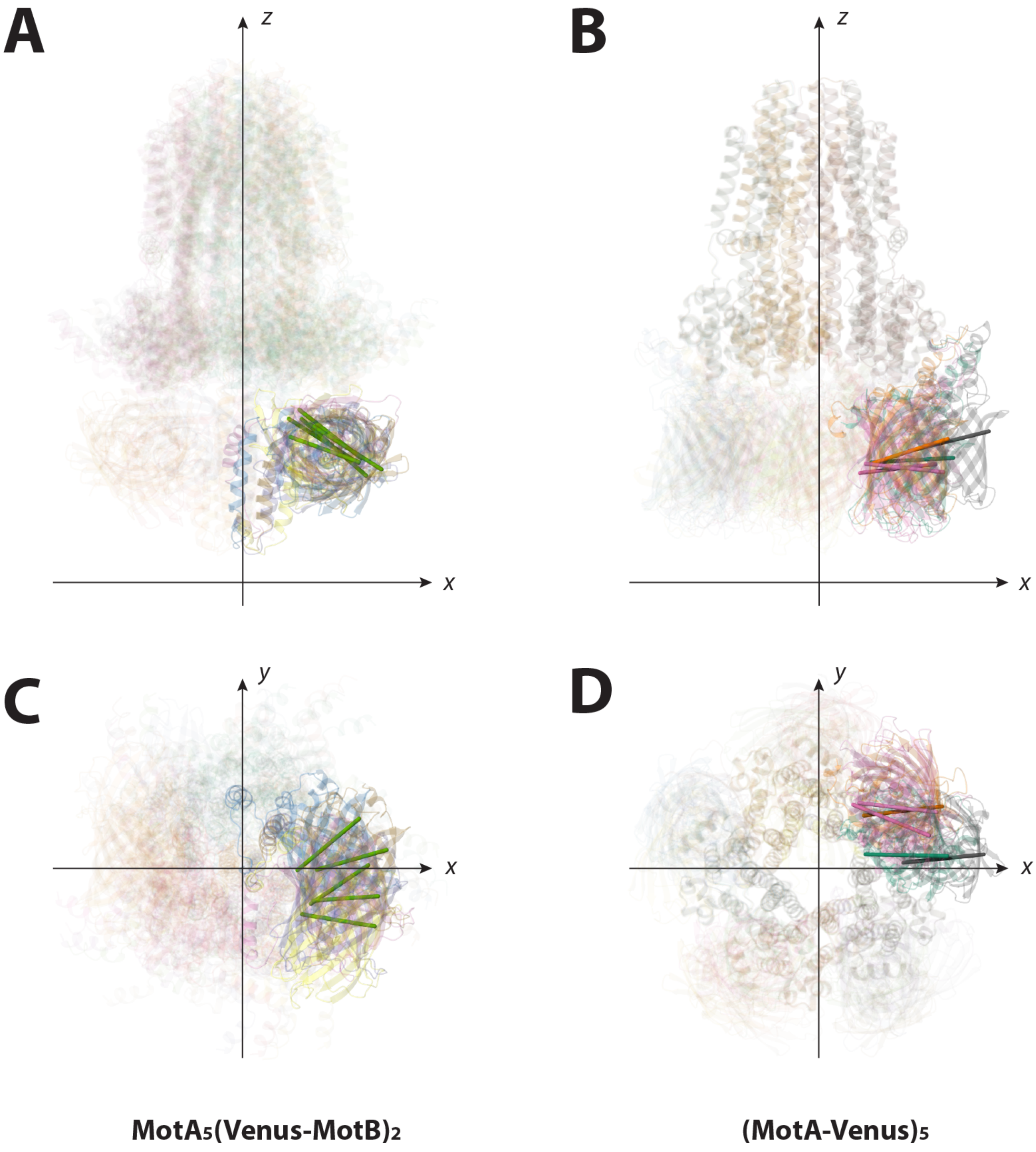


**Fig. S8. Fluorophore angular orientation distribution**. Approximations of the chromophore orientations (thick lines of various colors across the Venus barrels) in cartoon representations of predicted structures of different ranks. (**A**) Superimposed cartoon representations of the first five ranked predictions of $MotA_5(Venus\text{-}MotB)_2$, aligned by a peptide region spanning from the C-terminus end of GCN4 to the first 22 amino-acid residues at the N-terminus of MotB (assumed to provide a rigid reference). (**B**) Superimposed cartoon representations of the first five ranked predictions of $(MotA\text{-}Venus)_5$, aligned by one of the MotA chains in the $MotA_5$ ring. (**C**, **D**) Bottom views of the structures in the panels above.

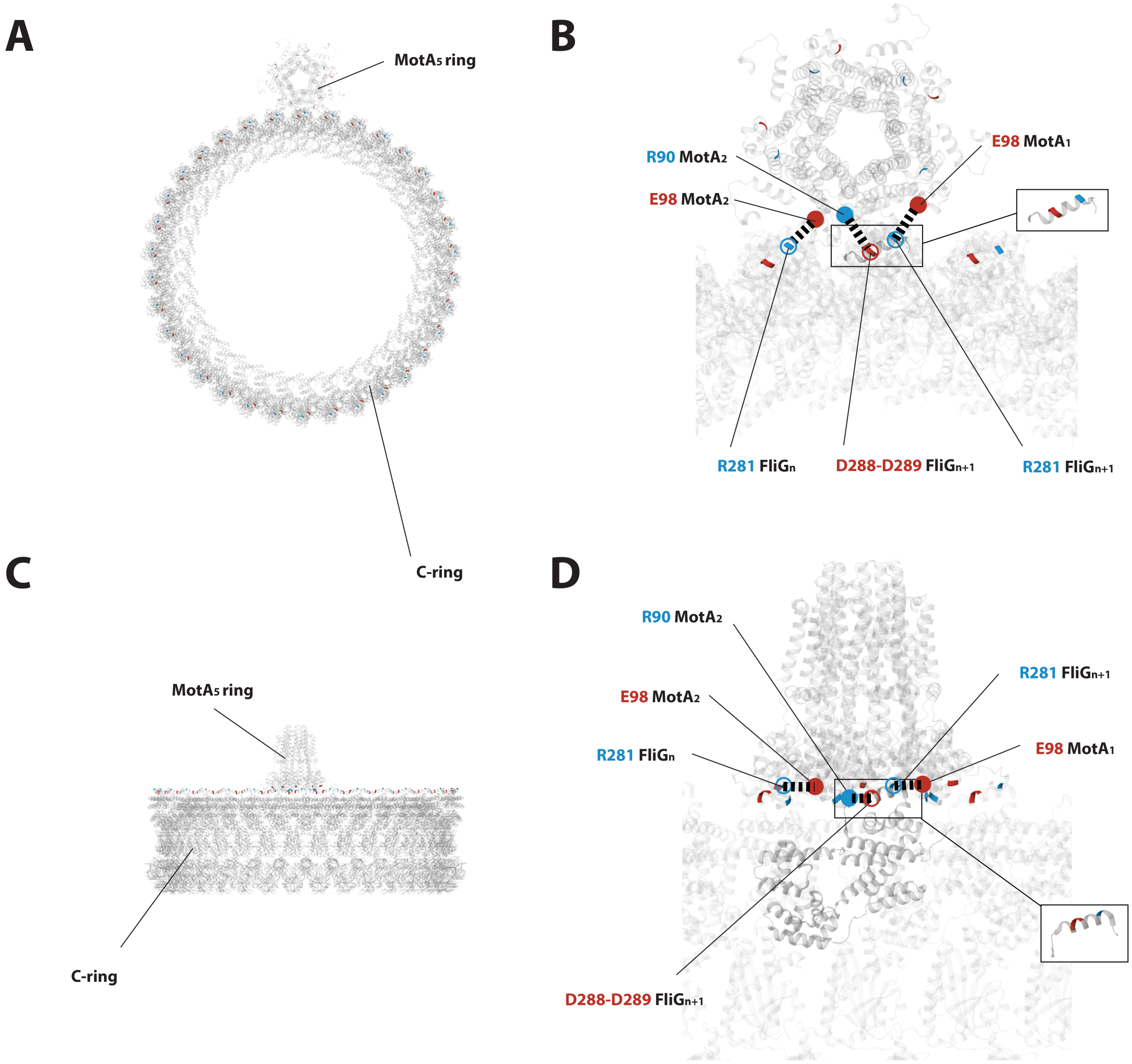


**Fig. S9. MotA-FliG interface in WT motors**. Cartoon representations of a $MotA_5$ stator ring, a C-ring in CCW conformation and their possible interaction. The spatial distribution of the charged residues in $MotA_5$ (emphasized by *filled red* and *blue circles*) and $FliG_{34}$ (emphasized by *open red* and *blue circles*) suggests a ***side-to-side*** interaction (see **MotA-FliG interaction** supporting text). Views form above (the extracellular space, **A**) and side (**C**). **B**: magnified view of the structures in **A**. Positively (blue) and negatively (red) charged amino acid residues in the FliG "torque coil" (inset) at the outer side of the C-ring interact electrostatically (*black dashed lines*) with complementary charged residues at the base of the $MotA_5$ ring. The FliG torque coil fits in the space between two adjacent MotA monomers in $MotA_5$. CW rotation of $MotA_5$ moves the C-ring to the left (CCW). **D**: Side view of the interactions depicted in **B**, as seen from the inside of the C-ring. The $MotA_5$ ring is an AlphaFold prediction of the *E. coli* MotA pentamer. The 34-symmetry CCW C-ring chosen for representation is the cryo-EM structure of *Salmonella enterica Typhimurium* C-ring (structure 8UOX on the RCSB Protein Data Bank). (3)

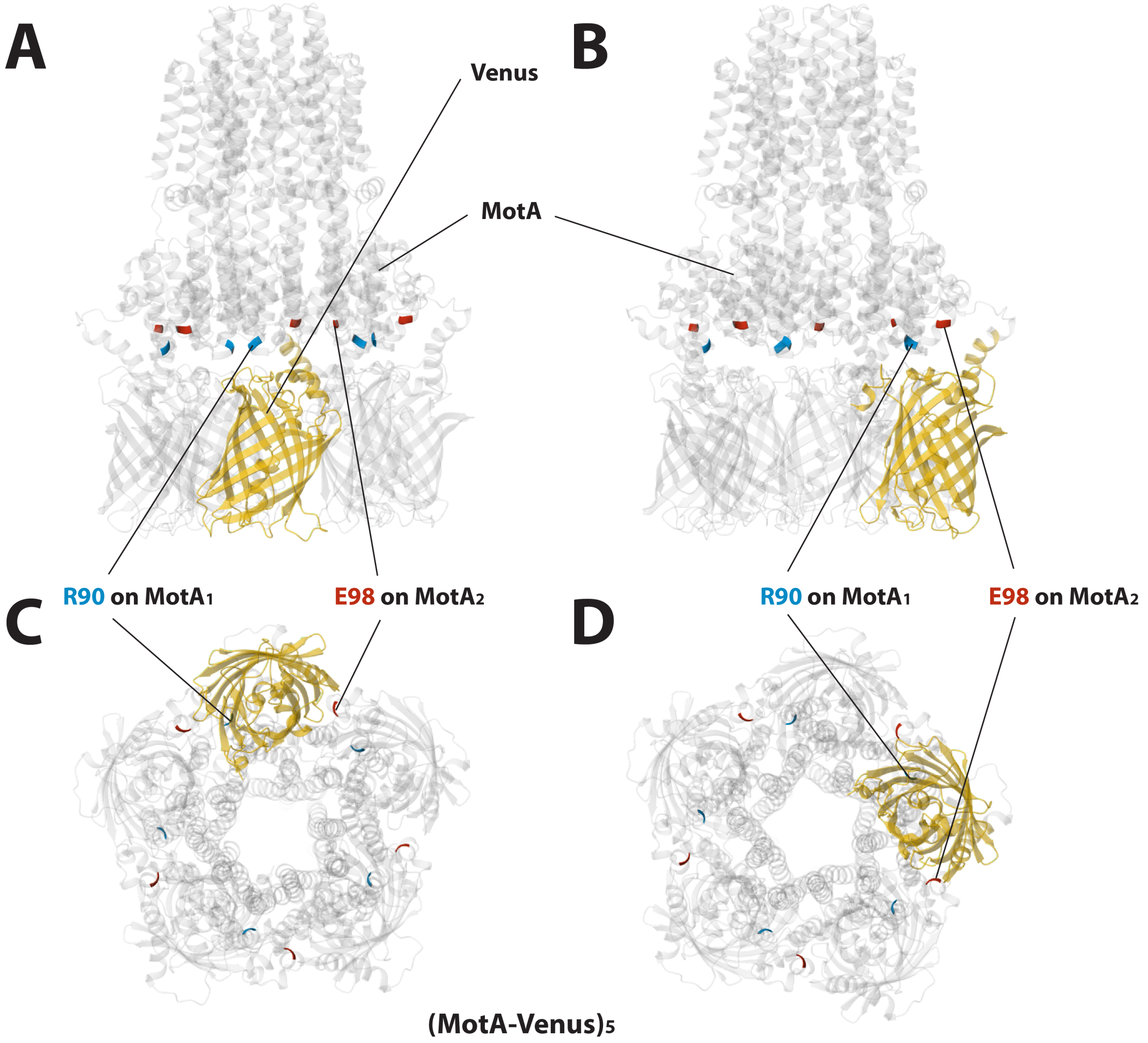


**Fig. S10. MotA-FliG interface in (MotA-Venus)$_5$ tagged motors**. Location of the fluorescent tags with respect to the site of interaction with FliG. Cartoon representations of (MotA-Venus)$_5$ viewed from the side (**A**, **B**), and from the bottom (**C**, **D**). Views in **B** and **D** are rotated with 90° along *z* axis. All charged residues involved in torque coupling are colored in *blue* (positive) or *red* (negative). The labeled ones interact with FliG. All Venus tags are located under the MotA$_5$ ring. A ***side-to-side*** interaction between the charged areas at the base of MotA$_5$ and at the upper-lateral side of the C-ring is possible. The fluorescence protein that tags MotA$_1$ (colored in *yellow*) does not obstruct the cog-wheel interface, but its linkage might. At the time of interaction with FliG, the Venus barrel might slide or pivot about flexible points in its linkage, possibly resulting in a periodic change of its orientation.

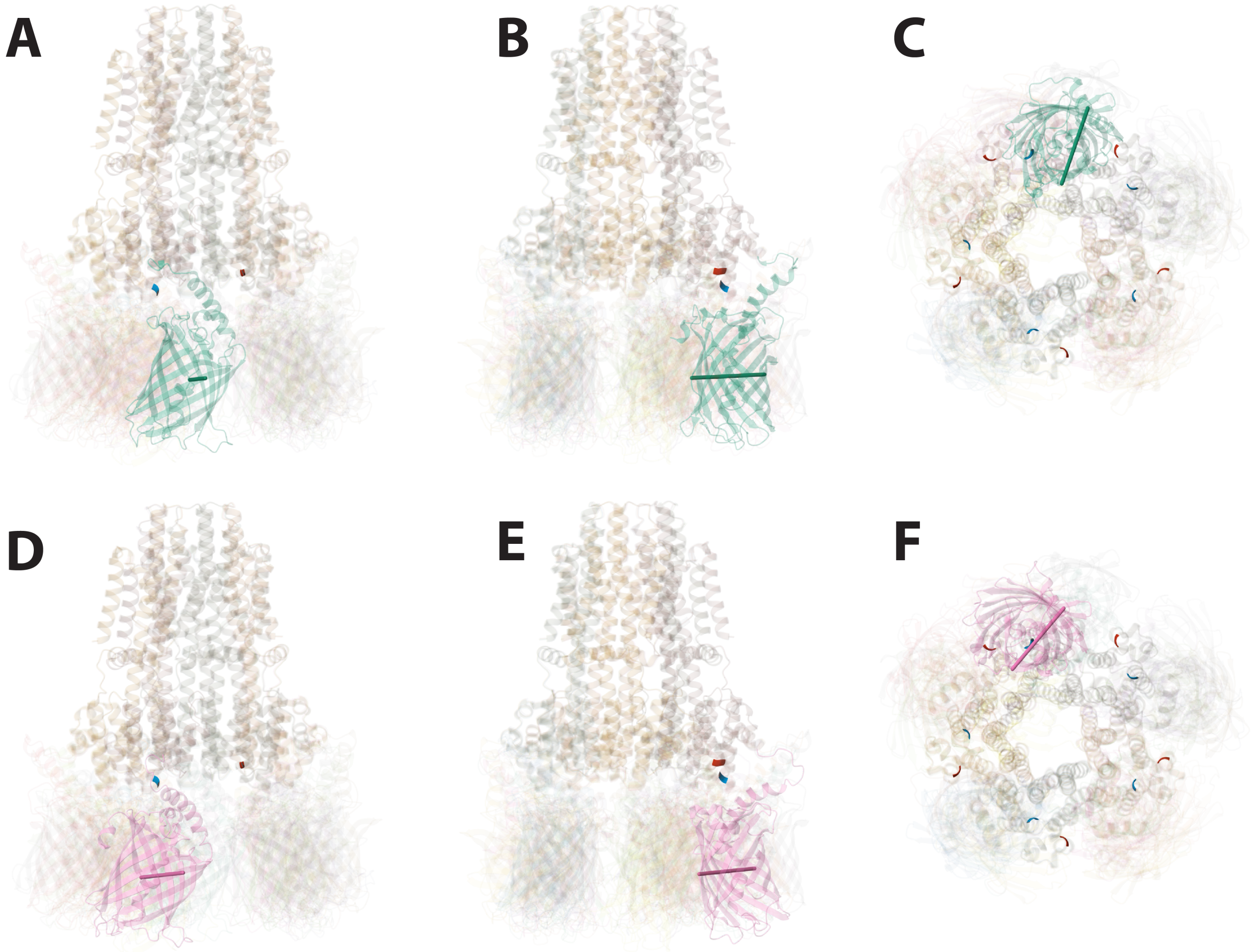


**Fig. S11. MotA-FliG interface in (MotA-Venus)$_5$ tagged motors**. Possible positioning of fluorescent tags with respect to the site of interaction with FliG in (MotA-Venus)$_5$ that allows the motor to function. Cartoon representations of the predicted structure in Fig. S10 viewed from the side (**A**, **B**) and below (**C**). **D-F**: alternative prediction of the same structure in which the linkage between the Venus tag and MotA does not obstruct the R90 charged residue in the MotA cleft above.

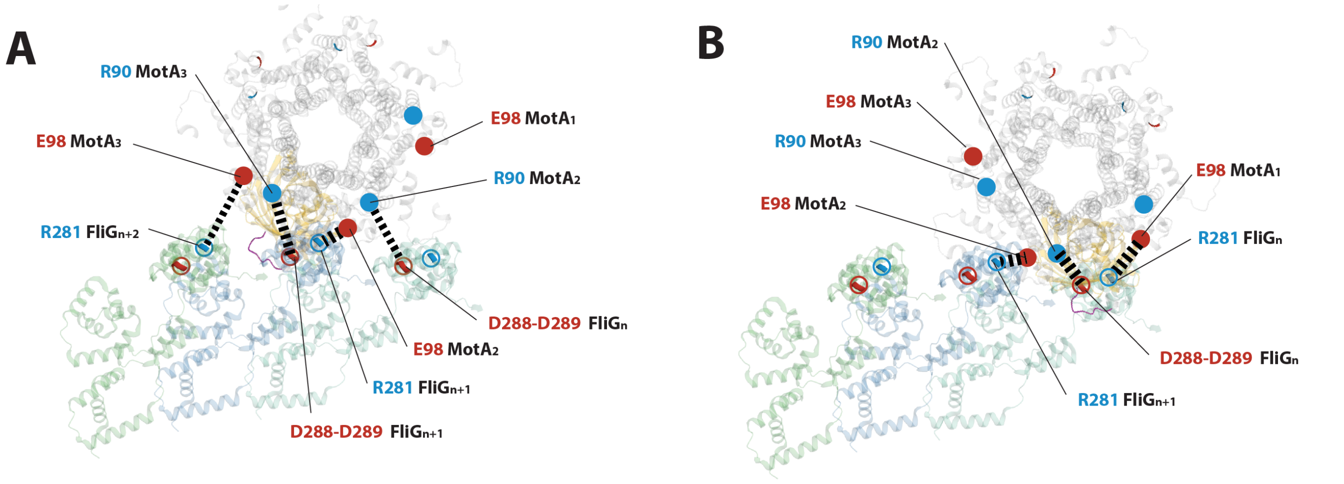


**Fig. S12. MotA-FliG interface in (Venus-MotA)$_5$ tagged motors**. Possible locations of the fluorescent tag (*yellow*) at the site of interaction with FliG in a side-to-side interaction MotA$_5$-FliG$_{34}$, consistent with a cog-wheel model. Two consecutive steps of MotA$_5$ (*light gray*), before (**A**) and after (**B**) a 36° CW rotation along a CCW C-ring. Views from the periplasmic space. *Black dashed lines*: electrostatic interactions between complementary charged residues on MotA (emphasizewd with filled red and blue circles) and FliG (emphasized with open red and blue open circles). Three adjacent FliG monomers (*light green* and *light blue*) of a 34-symmetry CCW *Salmonella enterica Typhimurium* C-ring (for representation, structure 8UOX on the RCSB Protein Data Bank (3)).

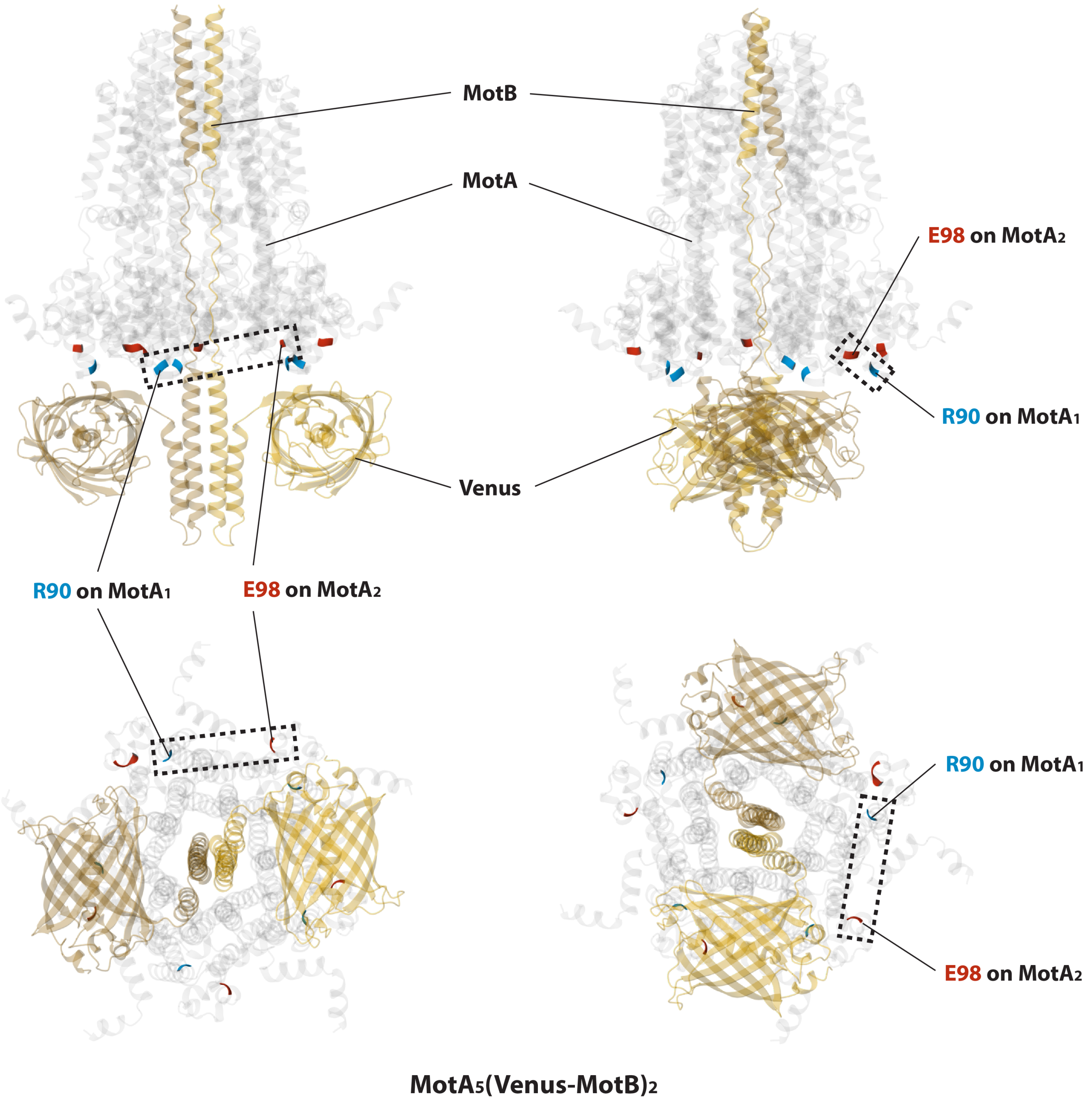


**Fig. S13. MotA-FliG interface in (Venus-MotB)$_2$ tagged motors**. Location of the fluorescent tags with respect to the site of interaction with FliG. Cartoon representations of (Venus-MotB)$_2$ viewed from the side (**A**, **B**), and from the bottom (**C**, **D**). Views in **B** and **D** are rotated with 90°along *z* axis. All charged residues involved in torque coupling are colored in *blue* (positive) or *red* (negative). A pair such as the boxed labeled ones interacts with FliG. Both Venus tags are located under the MotA$_5$ ring, allowing a side-to-side interaction with FliG$_{34}$.

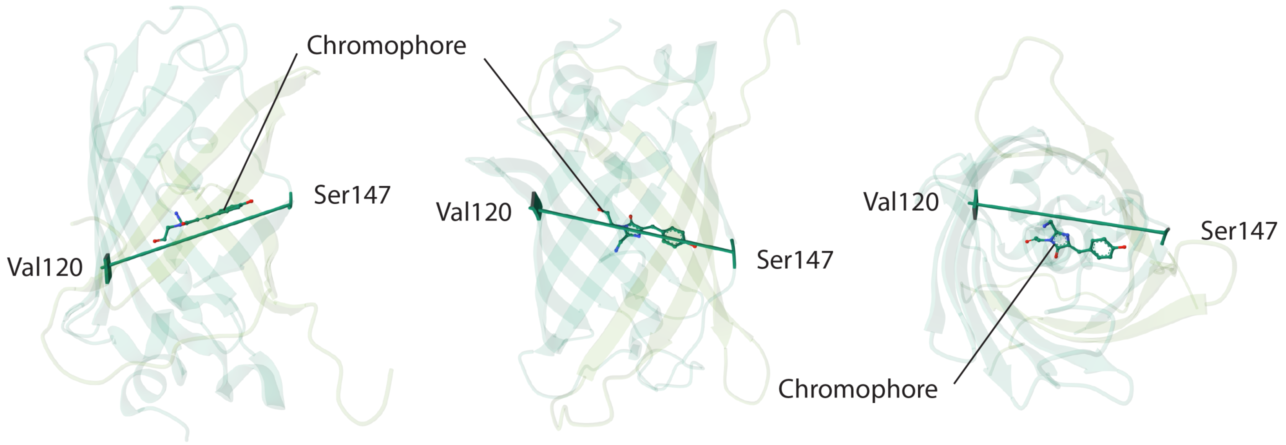


**Fig. S14. Approximation of chromophore orientation in a Venus fluorescent protein.**

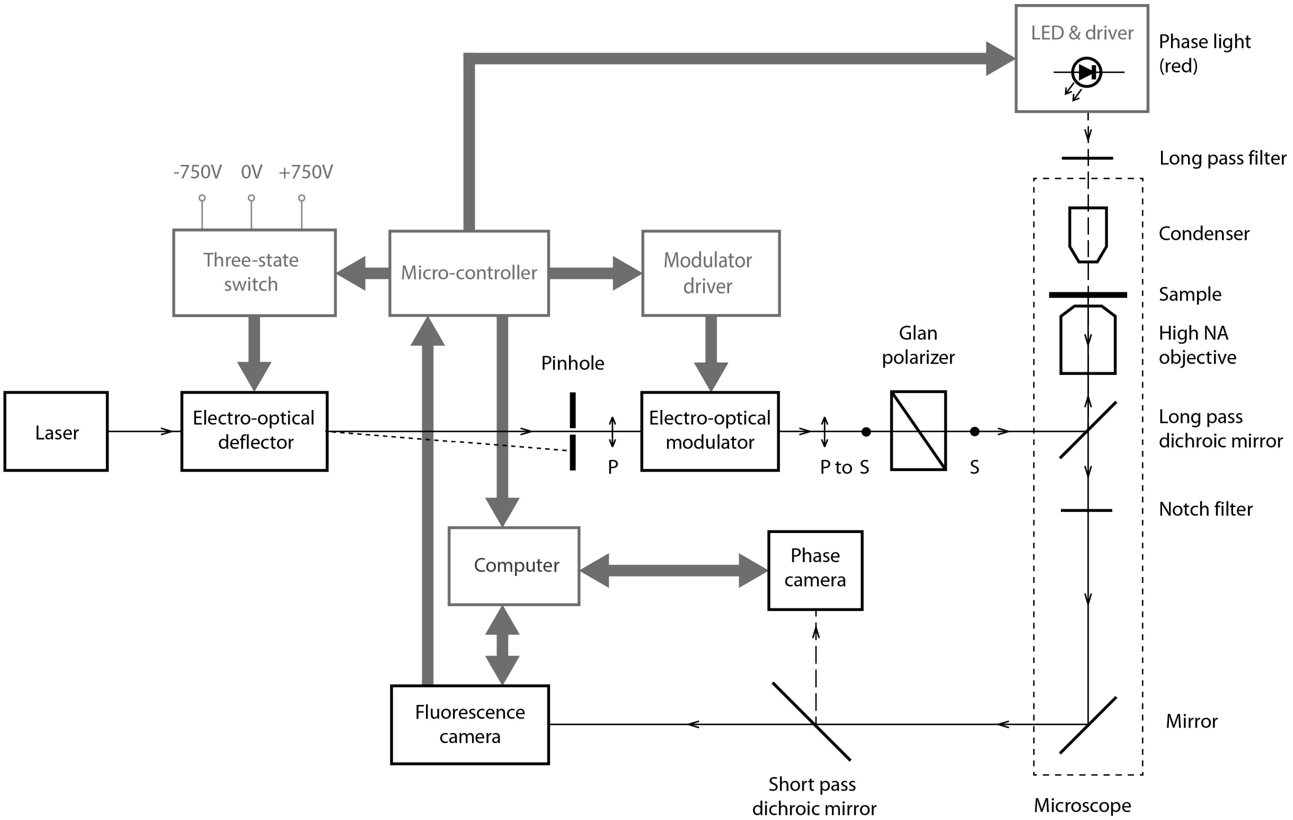


**Fig. S15. Schematic of Experimental Setup**. Fluorescence excitation laser light pulses of controlled duration (via an electro-optical deflector, which directs the laser beam through the center of a pinhole [ON], or away from it [OFF], as a function of the applied voltage) and intensity (via an electro-optical modulator, a voltage-controlled "variable" wave plate followed by a Glan polarizer) are applied to a modified inverted microscope. The excitation is directed to the input aperture of the objective through a long-pass dichroic mirror. Pulsed bright field illumination is provided by a red LED, via a condenser lens. The fluorescence emission and the bright field light are directed to the sensors of an EMCCD camera (for fluorescence or combined fluorescence/bright field imaging) and a CCD camera (for bright field imaging) via short pass dichroic mirror. A micro-controller triggers the computer video acquisitions of the two cameras, synchronizes the fluorescence excitation and phase light pulses with the corresponding camera frames and controls the amplitude and duration of the light pulses. A bleaching pulse of a longer duration and higher amplitude is applied at a prescribed time.

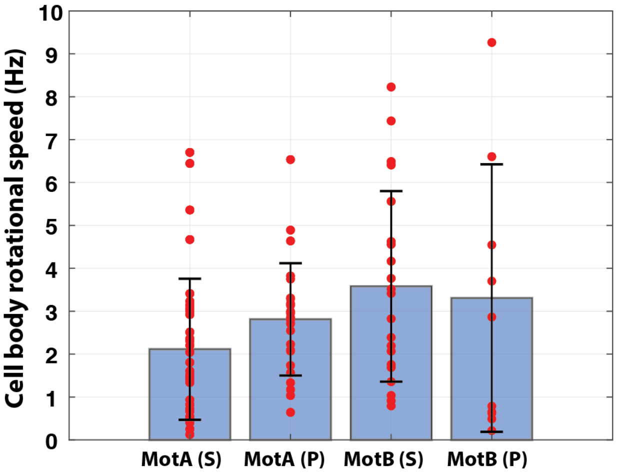


**Fig. S16. Rotational speed of tethered cells used in these experiments.** *Blue bars* with error bars: mean speed $\pm$ SD; *red filled circles*: individual tethered cell rotational speed values. From left to right: (MotA-Venus)$_5$ labeled cells with fluorescence excitation with S-polarized light: 2.1 $\pm$ 1.6 Hz (*n*=36), (MotA-Venus)$_5$ labeled cells with fluorescence excitation with P-polarized light: 2.8 $\pm$ 1.3 Hz (*n*=16), (Venus-MotB)$_2$ labeled cells with fluorescence excitation with S-polarized light: 3.6 $\pm$ 2.2 Hz (*n*=21) and (Venus-MotB)$_2$ labeled cells with fluorescence excitation with P-polarized light: 3.3 $\pm$ 3.1 Hz (*n*=11)

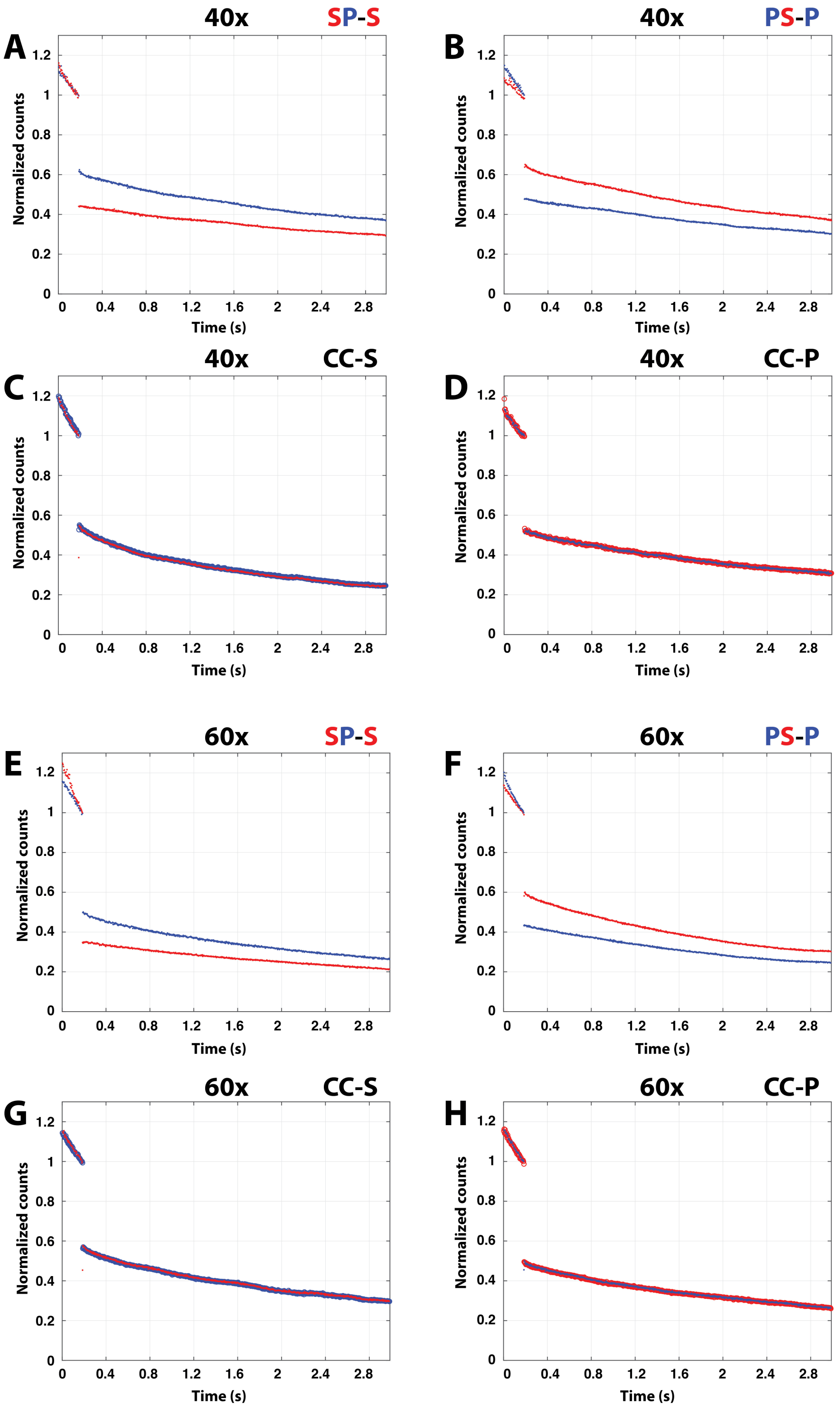


**Fig. S17. Experiments on beads.** Excitation using alternate pulses of S-polarized light (*red traces*) and P-polarized light (*blue traces*) (**A**, **B**, **E**, **F**) with a strong photo-bleaching pulse near 0.2 s (emission during bleaching pulse not represented). In all left panels, the photo-bleaching pulse was provided with S-polarized light. In all right panels, the photo-bleaching pulse was provided with P-polarized light. These measurements demonstrate polarization-specific photo-bleaching and polarization-dependent fluorescence emission with our setup. As expected, excitation with circularly polarized light (*red dots* and *blue circles*, each odd/even data point color-coded as in the panel above, see **Methods**), which does not preferentially photo-select fluorophores based on their orientation, resulted in similar fluorescence decay regardless of the polarization of the bleaching pulse (**C**, **D**, **G**, **H**). Experiments were performed with a 40x (epi-fluorescence mode, **A**-**D**) and 60x objective (TIRF mode, **E**-**H**).

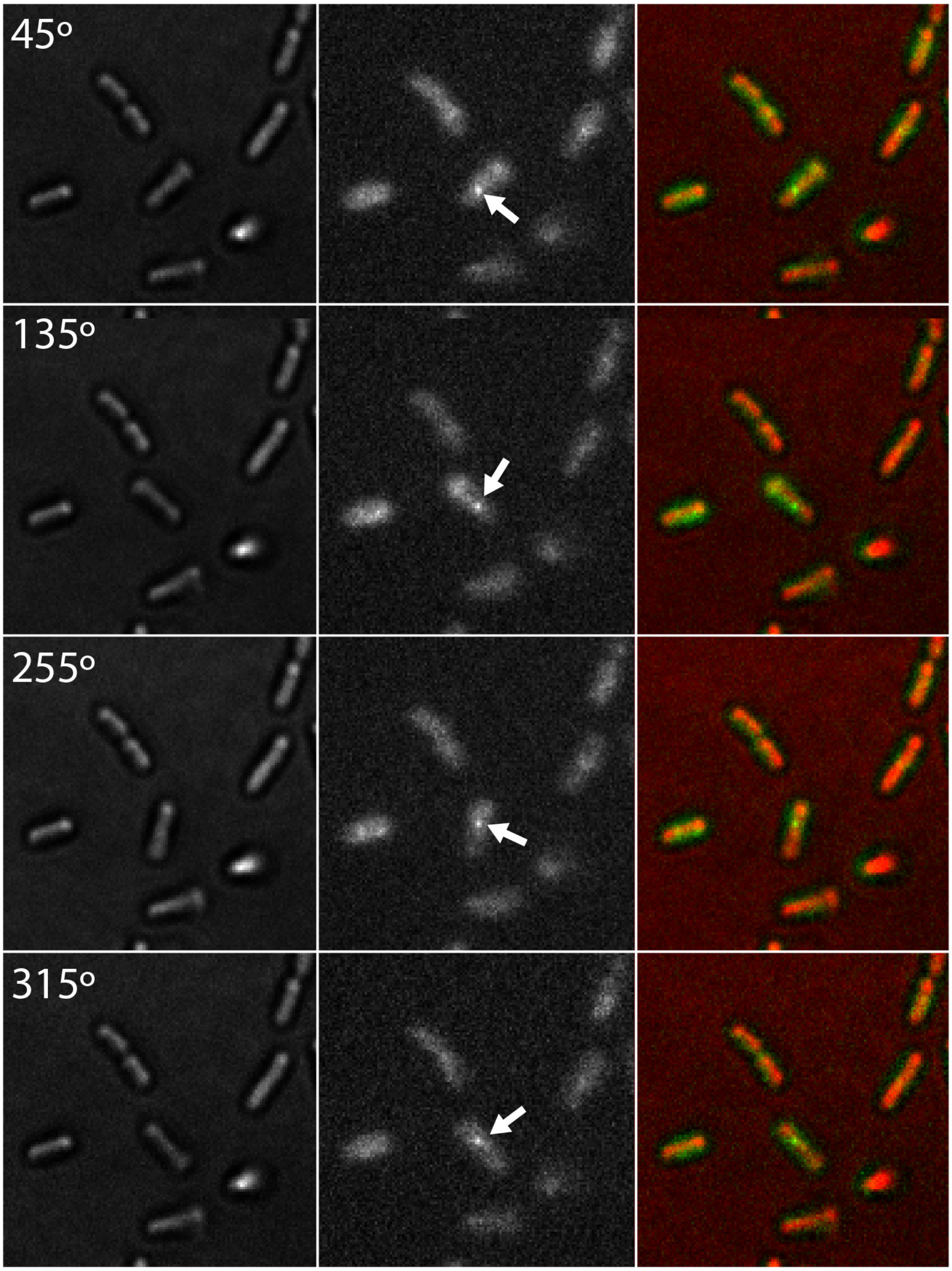


**Fig. S18. Tethered cell rotating around a MotA fluorescently labeled motor** (indicated by a white arrow). Synchronized bright field *(left column)* and fluorescence *(middle column)* acquisitions; *right column:* pseudo-colored bright field (red) and fluorescence (green) overlay for motor localization. Each image is the average of about 170 (bright field, 300 µs exposure) or 30 (fluorescence, 50 µs exposure) camera frames in which the cell body had the same orientation (from top to bottom row: 45°, 135°, 255° and 315°). See also Supplemental Movie S4.

**Table S1. Average fluorescence emission intensities of the motor spots of tethered cells with MotA-Venus and Venus-MotB labeled motors**

| | **Excitation in S** | **Excitation in P (control)** |
|---|---|---|
| $(\text{MotA-Venus})_5$ fluorescence intensity (Counts, mean $\pm$ SD) | 3045.8 $\pm$ 1790.3 ($n$=22) | 2419.2 $\pm$ 1251.1 ($n$=27) |
| $(\text{Venus-MotB})_2$ fluorescence intensity (Counts, mean $\pm$ SD) | 859.6 $\pm$ 115.6 ($n$=24) | 866.9 $\pm$ 303.1 ($n$=8) |
| $(\text{MotA-Venus})_5$ : $(\text{MotA-Venus})_2$ mean fluorescence intensity ratio (times) | 3.4 : 1 | 2.8 : 1 |
| Bootstrap $(\text{MotA-Venus})_2$:$(\text{MotB-Venus})_2$ ratio error estimation (times $\pm$ SD) | 3.34 $\pm$ 0.41 : 1 $\pm$ 0.09 | 2.82 $\pm$ 0.27 : 1$\pm$ 0.13 |

Data represents the average fluorescence emission intensities of all motors used in this study, as acquired in the first fluorescence frame. Only the motors probed under identical acquisition conditions were included: TIRF mode, fluorescence excitation time (set to 50 µs) and camera electron multiplying (EMCCD) factor (set to 100). To increase the number of samples, we included data from experiments discarded for the reason of switching the direction of rotation during acquisition but met the other criteria (rotation centered to a bright spot and significant step-wise fluorescence bleaching after the bleaching pulse was applied).

**Table S2. Sequences of engineered proteins**

| Name | Sequence |
|---|---|
| Venus-MotB | MSKGEELFTGVVPILVELDGDVNGHKFSVSGEGEGDATYGKLTLKLLCTT<br>GKLPVPWPTLVTTLGYGVQCFARYPDHMKQHDFFKSAMPEGYVQERTIFF<br>KDDGNYKTRAEVKFEGDTLVNRIELKGIDFKEDGNILGHKLEYNYNSHNV<br>YITADKQKNGIKANFKIRHNIEDGGVQLADHYQQNTPIGDGPVLLPDNHYL<br>SYQSKLFKDPNEKRDHMVLLEFLTAAGITHGEGEAQKQKEQRQAAEELAN<br>GSGGSKQLEDKVEELLSKNYHLENEVARLKKLVGERPIIVVKRRKAKSHG<br>AAHGSWKIAYADFMTAMMAFFLVMWLISISSPKELIQIAEYFRTPLATAV<br>TGGDRISNSESPIPGGGDDYTQSQGEVNKQPNIEELKKRMEQSRLRKLRG<br>DLDQLIESDPKLRALRPHLKIDLVQEGLRIQIIDSQNRPMFRTGSADVEP<br>YMRDILRAIAPVLNGIPNRISLSGHTDDFPYASGEKGYSNWELSADRANA<br>SRRELMVGGLDSGKVLRVVGMAATMRLSDRGPDDAVNRRISLLVLNKQAE<br>QAILHENAESQNEPVSALEKPEVAPQVSVPTMPSAEPR |
| MotA-Venus | VLILLGYLVVLGTVFGGYLMTGGSLGALYQPAELVIIAGAGIGSFIVGNNGKA<br>IKGTLKALPLLFRRSKYTKAMYMDLLALLYRLMAKSRQMGMFSLERDIENPR<br>ESEIFASYPRILADSVMLDFIVDYLRLIISGHMNTFEIEALMDEEIETHES<br>EAEVPANSLALVGDSLPAFGIVAAVMGVVHALGSADRPAAELGALIAHAMV<br>GTFLGILLAYGFISPLATVLRQKSAETSKMMQCVKVTLLSNLNGYAPPIAVEF<br>GRKTLYSSERPSFIELEEHVRAVKNPQQQTTTEEAAKEAAAKEAASKGEELFT<br>GVVPILVELDGDVNGHKFSVSGEGEGDATYGKLTLKLICTTGKLPVPWPTLV<br>TTLGYGVQCFARYPDHMKQHDFFKSAMPEGYVQERTIFFKDDGNYKTRAEV<br>KFEGDTLVNRIELKGIDFKEDGNILGHKLEYNYNSHNVYITADKQKNGIKANF<br>KIRHNIEDGGVQLADHYQQNTPIGDGPVLLPDNHYLSYQSKLSKDPNEKRDH<br>MVLLEFVTAAGITHGMDELYK |
| Sticky FlgE | MAFSQAVSGLNAAATNLDVIGNNIANSATYGFKSGTASFADMFAGSKVGLGV<br>KVAGITQDFTDGTTTNTGRGLDVAISQNGFFRLVDSNGSVFYSRNGQFKLDE<br>NRNLVNMQGLQLTGYPATGTPPTIQQGANPTNISIPNTLMAAKTTTTASMQI<br>NLNSSDPLPTVTPFSASNADSYNKKGSVTVFDSQGNAHDMSVYFVKTGDNNW<br>QVYTQDSSDPNSIGLNRQSFRGRWHEAKTATTLEFNANGTLVDGAMANNIAT<br>GAINGAEPATFSLSFLNSMQQNTGANNIVATTQNGYKPGDLVSYQINDDGTV<br>VGNYSNEQTQLLGQIVLANFANNEGLASEGDNVWSATQSSGVALLGTAGTGN<br>FGTLTNGALEASNVDLSKELVNMIVAQRNYQSNAQTIKTQDQILNTLVNLR |

**Movie S1. AlphaFold prediction of Venus barrel orientations in (Venus-MotB)$_2$. Only the periplasmic part of MotB is shown. Link to Movie.**

**Movie S2. AlphaFold prediction of Venus barrel orientations in (MotA-Venus)$_2$, within MotA$_5$. Link to Movie.**

**Movie S3. Chromophore TDM approximation in Venus)$_5$. Link to Movie.**

**Movie S4. Tethered cell rotating around a bright motor spot, probed simultaneously in bright field (left panel) and fluorescence microscopy (middle panel). The right panel is a pseudo-colored overlay of the bright field (red) and fluorescence (green) frames, for co-localization. Each frame in the movie is the average all acquired frames in which the cell had the same orientation, in 30° bins. Link to Movie.**
